# *Meteora sporadica*, a protist with incredible cell architecture, is related to Hemimastigophora

**DOI:** 10.1101/2023.08.13.553137

**Authors:** Yana Eglit, Takashi Shiratori, Jon Jerlström-Hultqvist, Kelsey Williamson, Andrew J. Roger, Ken-Ichiro Ishida, Alastair G. B. Simpson

## Abstract

‘Kingdom-level’ branches are being added to the tree of eukaryotes at a rate approaching one per year, with no signs of slowing down^1–4^. Some are completely new discoveries, while others are morphologically unusual protists that were previously described but lacked molecular data. For example, Hemimastigophora are predatory protists with two rows of flagella that were known since the 19^th^ century, but proved to represent a new deep-branching eukaryote lineage when phylogenomic analyses were conducted^2^. *Meteora sporadica* Hausmann et al. 2002^5^ is a protist with a unique morphology and motility; cells glide over substrates along a long axis of anterior and posterior projections, and have a pair of lateral ‘arms’ that swing back and forth. Originally, *Meteora* was described by light microscopy only, from a short-term enrichment of deep-sea sediment. A small subunit ribosomal RNA (SSU rRNA) sequence was reported recently, but the phylogenetic placement of *Meteora* remained unresolved^6^. Here, we investigated two cultivated *Meteora sporadica* isolates in detail. Transmission electron microscopy showed that the anterior-posterior projections are supported by microtubules originating from a cluster of subnuclear MTOCs. Likewise, the arms are supported by microtubules, and neither have a flagellar axoneme-like structure. Sequencing the mitochondrial genome showed this to be amongst the most gene-rich known, outside jakobids. Remarkably, phylogenomic analyses of 254 nuclear protein-coding genes robustly support a close relationship with Hemimastigophora. Our study suggests that *Meteora* and Hemimastigophora together represent a morphologically diverse ‘supergroup’, and thus are important for resolving the tree of eukaryote life and early eukaryote evolution.

## Results and Discussion

### Morphology

The two *Meteora sporadica* isolates, SRT610 (Fig 1A-B) and LBC3 (Fig 1C-F; Fig S1A-F), have a similar morphology. The cell body is 4.4 ± 0.6 μm long and 3.6 ± 0.4 μm wide in isolate SRT610 (n=22) and 4.3 ± 0.9 μm by 3.2 +/- 0.7 μm in isolate LBC3 (n=24). The anterior projection is 8.1 ± 1.7 μm long in SRT610 (n=22) and 13.0 ± 4.5 μm in LBC3 (n=24), while the posterior is 6.7 ± 1.6 μm and 8.4 ± 2.9 μm, respectively. There are typically two lateral ‘arms’ of length 2.7 ± 0.5 μm or 2.6 ± 0.9 μm emerging from the cell body, but some individuals have more (Fig 1D, Supplementary Video 2). The cell glides along the surface via its long axis (Supplementary Video 1). The arms normally swing regularly back and forth, but gliding persists when the arms are static or absent (Supplementary Video 3), indicating that this motility does not depend on arm movement. Detached floating cells bend and squirm, but appear to lack directed motility. There are numerous small granules along the arms, as well as the long axis (Fig 1), most of which correspond to extrusomes (see below). These granules move back and forth along both arms and the long axis (Supplementary Video 1), as well as between them (Fig S1D). Occasionally, protrusions up to 3 μm long can extend rapidly from both arms and the long axis (Fig S1D).

**Figure 1.**
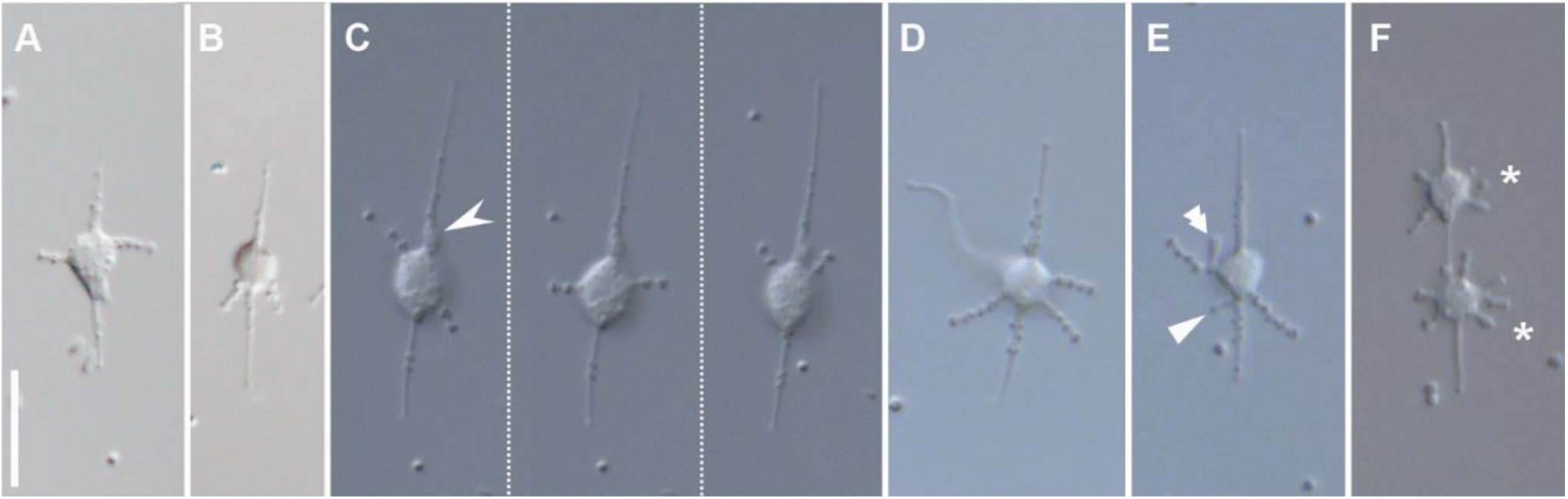
DIC light micrographs of *Meteora sporadica* isolates SRT610 (A-B) and LBC3 (C-F). A-B) General views of two individuals. C) One cell at different time points showing typical positions of lateral ‘arms’. A partially ingested bacterium can be seen anterior of the cell body proper (barbed arrowhead). D) Individual with three lateral arms and a large cytoplasmic extension (arrow). E) Individual with two small extensions, one from the long axis (single arrowhead) and another from a lateral arm (double arrowhead). More images of this cell in Fig S1D. F) Late cell division, with two individuals (asterisks) separating along the longitudinal axis. Scale bar: A-F (in A) 10 µm.

Transmission electron microscopy (TEM) examination of the cell body shows the vesicular nucleus, food vacuoles containing bacteria, and mitochondria (or possibly one ramified mitochondrion) located to the dorsal side of the nucleus (Fig 2A). Mitochondria have flat cristae (Fig S2B). Adjacent to the ventral side of the nucleus (Fig 2A, Fig S2C) is a flat cluster of about seven (5-8) microtubule organising centres (MTOCs), which are hexagonally arranged, mostly as two staggered rows (Fig 2B). Each MTOC is a cylindrical structure approximately 160 nm long (mean 158 ± 18 nm; n=9) and 120 nm wide (116 ± 56 nm; n=13) with a dense-staining core 100 nm long (100 ± 13 nm) and 55 nm (56 ± 7 nm) wide (Fig 2A, F, G; Fig S2E, M). At least some MTOCs are anchored to the nuclear envelope (Fig S2 C, D). Emerging microtubules form both longitudinal and transverse bundles (Fig 2B; Fig S2E, G, L-M). The arrangement of microtubules in longitudinal bundles is irregular in cross section (Fig 2C). Transverse microtubules converge at the base of arms, then extend into them (Fig S2G). The cell body, longitudinal extensions and arms contain numerous ovoid extrusomes approximately 240 nm long (243 ± 16 nm; n=11) and 180 nm wide (178 ± 20 nm; n=11), composed of mostly of light-staining material but with a dark staining cylinder at the base (Fig 2D). Some vacuoles, including food vacuoles, are coated by fibrillar material on the inside (Fig 2E).

**Figure 2.**
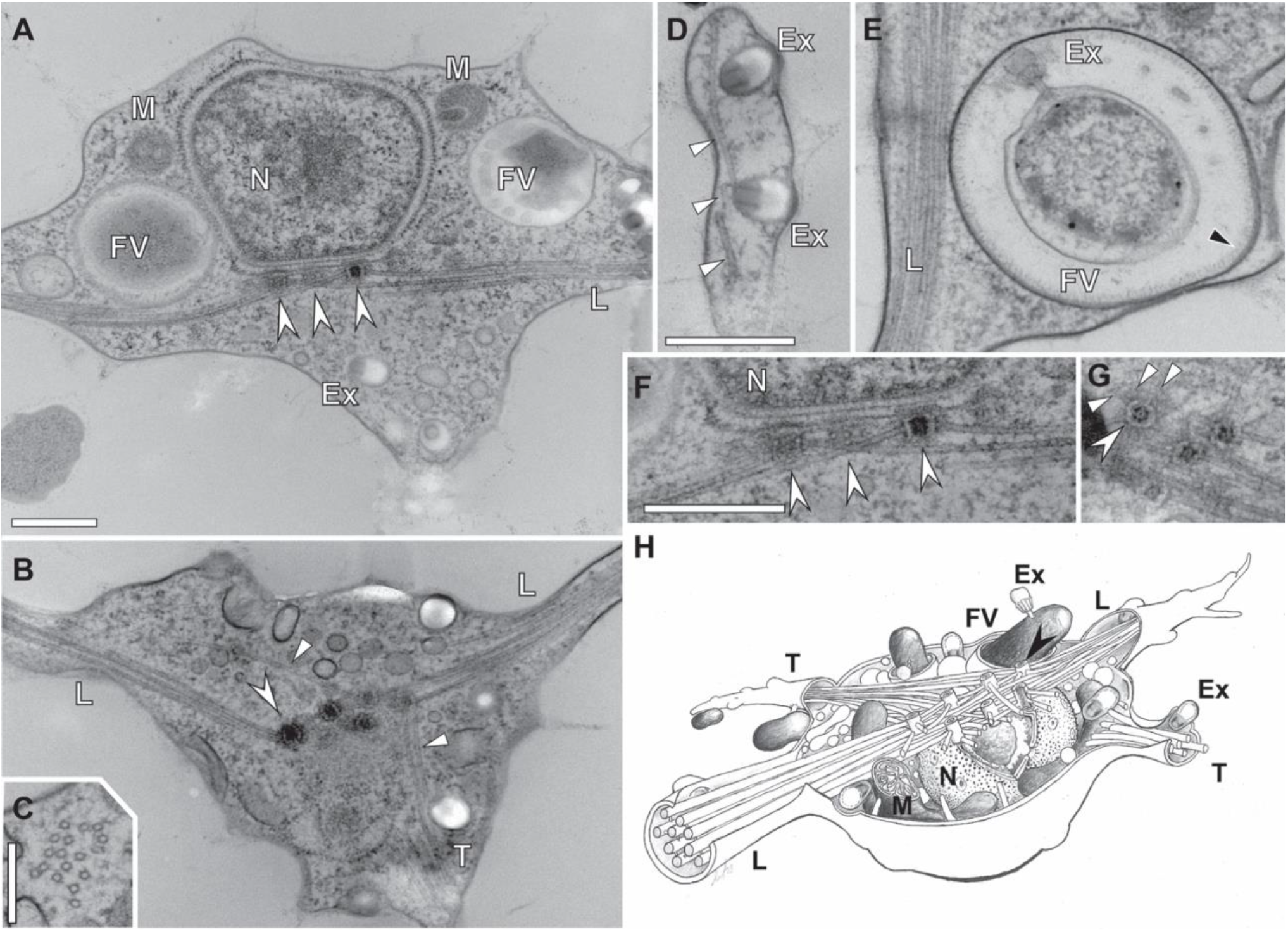
Ultrastructure of *Meteora sporadica* isolate SRT610 as imaged by transmission electron microscopy. A) Section through the cell body with the dorsal side to top of image. A cluster of microtubule organising centres (MTOCs, barbed arrowheads) is attached on the ventral surface of the nucleus (N). The longitudinal bundle of microtubules (L) passes to the left and right from the MTOCs. Mitochondrial sections (M) and food vacuoles (FV) can be seen. B) Cross section of the cell body and cluster of MTOCs (barbed arrowhead), longitudinal bundle (L) extending to the left and right of the image. Transverse microtubules (T, arrowhead) emerge from the MTOCs and are seen here extending to the bottom of the image. For the adjacent section in the series of the same cell, see Fig S2F. C) Cross section through the longitudinal bundle of microtubules. D) Section through part of a lateral arm showing microtubules (arrowheads) and two extrusomes (Ex). E) Discharged extrusome (Ex) attached to the surface of a prey bacterium inside a food vacuole (FV). Note coating inside the vesicle (black arrowhead), and the longitudinal bundle (L). More images of discharged extrusomes can be seen in FigS2I-K. F) Detail of MTOC (barbed arrowheads) attachment to the nucleus (N), and emerging longitudinal microtubules, from Fig 2A. G) Cross-section through four MTOCs showing a radial emergence of microtubules (arrowheads). H) Diagram of the cell structure. Scale bars: A-B (in A) 500 nm, C) 250 nm, D-E (in D) 500 nm, F-G (in F) 500 nm.

*Meteora* feeds by contacting bacterial prey with an extrusome (Fig S1C, Supplementary Video 4), typically on one of the arms. The bacterium becomes attached, is gradually moved to the base of the arm, then is phagocytosed once at the cell body proper. A structure inferred to be a discharged extrusome was observed by TEM in most vacuoles with discernible prey material (Fig 2E, Fig S2I-K). Most microbial eukaryotes with extrusomes use them for capturing eukaryotic prey or for defense^7,8^; it is notable that *Meteora* uses extrusomes to capture prokaryotes, which is much rarer^9^.

The cells divide across the long axis (Fig 1F): the cell stops in place while the cell body proper moves slightly up and down for several minutes (Fig S1E) until visible cytokinesis begins (Fig S1F); the daughter cells pinch off and each re-establishes the missing end of the long axis in approximately 5 minutes.

Most unicellular organisms that glide across surfaces, eating bacteria, are flagellates, and these often glide on one of their flagella. They are highly abundant and found widely distributed across the tree of eukaryotes: examples include phagotrophic euglenids^10^, glissomonads^11^, mantamonads^12^, and apusomonads^13^. The ecology and behaviour of *Meteora* closely resembles that of bacterivorous gliding flagellates; however, *Meteora* does not have flagella, nor obvious derivatives. Thus, it defies assignment to one of the general ecological categories of eukaryotic microorganisms.

### rRNA analyses

An SSU rRNA gene phylogeny of 192 taxa broadly representing eukaryote diversity (Fig S3A) agreed with a similar recent analysis ^6^ in failing to resolve the phylogenetic position of *Meteora*. A similarly broad dataset of concatenated SSU+LSU rDNA likewise did not resolve the placement of *Meteora* with any support, nor place it in any major group of eukaryotes (Fig S3B).

A phylogenetic placement analysis of environmental sequences from publicly available datasets (see methods) using RAxML-EPA identified almost no candidate relatives of *Meteora*. This analysis assigned a likelihood-weight ratio of 0.7 to a marine environmental sequence (asv_053_06994, Biomarks), however this in turn is 97% identical to uncultured marine hydrothermal vent sediment clone AT4-68 (AF530543.1); this latter sequence sometimes resolves as sister to *Meteora* in SSU rRNA gene phylogenies, but without support (e.g. 20% bootstrap support in our analysis). A sequence from a neotropical soils metatranscriptome ^14^ was identified with a likelihood-weight ratio of only 0.56 and with just 86% sequence identity to *Meteora sporadica* (LBC3) – thus, its identity remains inconclusive. No other environmental sequence hits were found. Thus, *Meteora* appears to represent a distinct phylogenetic entity in the current molecular tree of eukaryotes.

### Phylogenomics

To better place *Meteora* in the eukaryote tree of life, we generated a transcriptome from each of our isolates. We then assembled a 254-gene dataset, representing a broad sampling of eukaryotic diversity through 108 taxa, reduced to 66 for computationally-intensive analyses. The phylogenomic marker genes were well represented in the sequenced transcriptomes (236/254 genes for both), which additionally had relatively high BUSCO scores (231 and 211 complete/255, SRT610 and LBC3, respectively; see methods). Phylogenies inferred for both the 108- and 66-taxon datasets broadly agree with other eukaryote-wide phylogenetic studies^1,2,4,15^, for example recovering Sar, Obazoa, Amorphea (i.e. Obazoa+Amoebozoa) and Discoba with full support (Fig 3, Fig S4A). We did not, however, recover Telonemia as the sister group to Sar (i.e. the TSAR group)^15^. Remarkably, *Meteora* did not fall into any of the well-established supergroups, but instead formed a maximally-supported clade with Hemimastigophora, a phylogenetically isolated taxon recently-proposed to represent a new eukaryote supergroup^2^. The heterotrophic flagellate *Ancoracysta* (representing a different newly proposed supergroup, Provora^16^) branches as sister to this *Meteora*-Hemimastigophora clade, though with weaker support (85% PMSF bootstrap support; 97% UFBOOT support; posterior probability 0.99)

**Figure 3.**
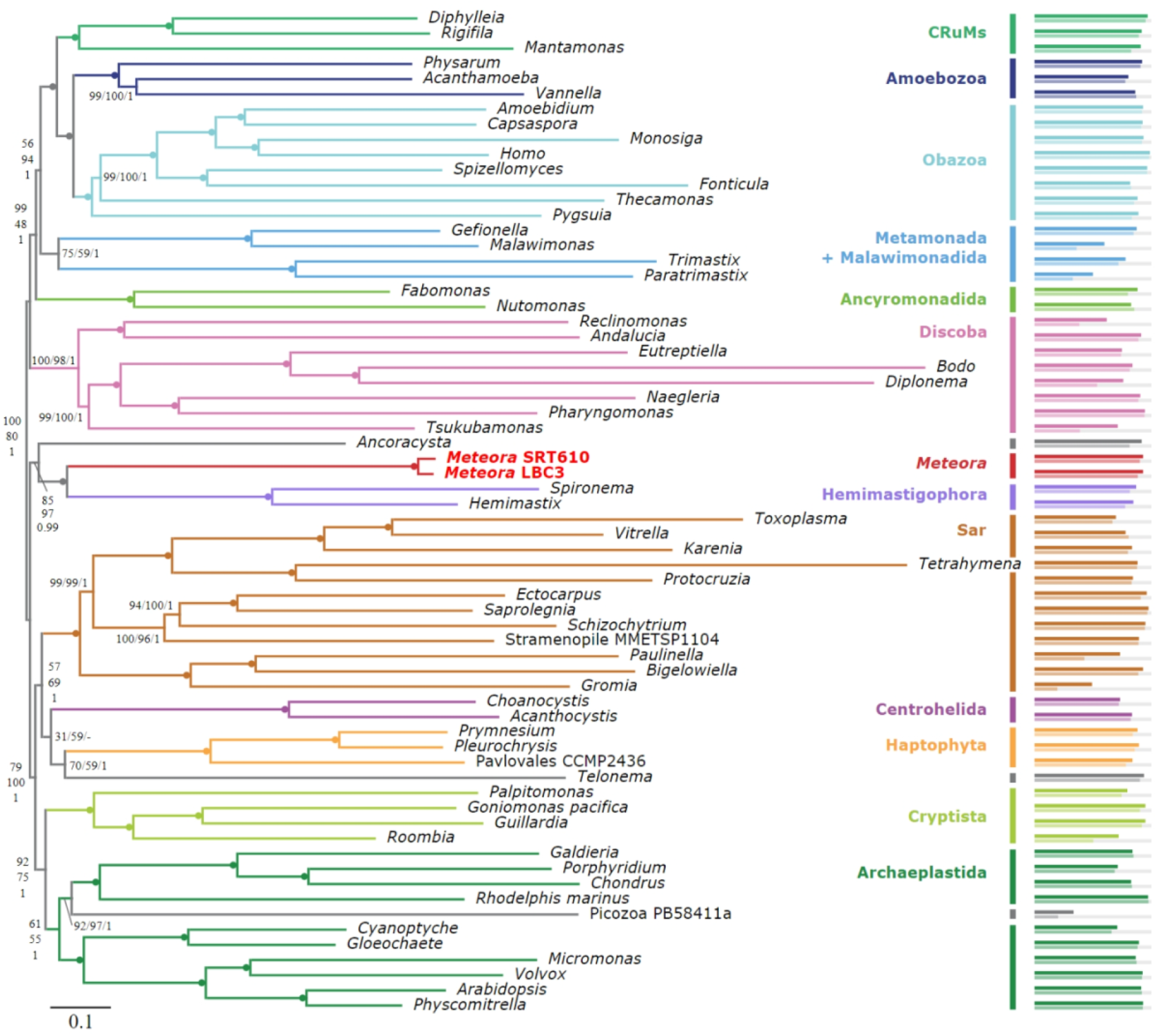
Phylogenetic placement of *Meteora* among eukaryotes. Maximum likelihood phylogeny inferred from 70471 sites across 254 genes over 66 taxa under the LG+MAM60+Γ model. Support values on branches show posterior mean site frequency bootstrap support (PMSF; 200 true replicates), UFBOOT support (1000 replicates), and Bayesian posterior probabilities (PP) under the CAT+GTR model, in that order, left to right or top to bottom. Filled circles indicate full support (100%, 100%, 1). Bars on the right indicate % coverage by gene (above) and by site (below).

To validate the robustness of the *Meteora*+Hemimastigophora clade, we examined multiple variations on the 66-taxon dataset that test for potential sources of phylogenetic error. Analysis of a dataset that excluded three branches identified as long-branching outliers (“nLB”) still returned maximal support for the *Meteora*+Hemimastigophora clade (Fig S4B). Recoding the amino acid data into a reduced alphabet of 4 classes based on (i) the pre-defined “SR4” categorisation ^17^, or (ii) custom classes optimised to minimise across-taxon compositional bias in this dataset (minmax-chisq)^18^, both robustly supported *Meteora*+Hemimastigophora (SR4 and MinMaxChisq: 100% and 99% UFBOOT support, respectively; Fig S4C-D). By contrast, these same analyses did not recover *Ancoracysta*+*Meteora*+Hemimastigophora and placed *Ancoracysta* elsewhere entirely, as sister to ‘Diaphoretickes’ (SR4, 92% UFBOOT) or to haptophytes (minmax-chisq, 97% UFBOOT). Incidentally, the 66-taxon dataset without *Ancoracysta* (noAnco) recovered the *Meteora*+Hemimastigophora relationship with full support (Fig S4E).

Removal of the fastest evolving sites in 10% increments (FSR analysis; Fig. S4G) showed *Meteora*+Hemimastigophora as maximally supported until 30% sites remaining, whereupon support dropped to ∼80% UFBOOT; the widely-accepted Discoba and CRuMs clades behave similarly. Conversely, support for *Ancoracysta*+*Meteora*+Hemimastigophora was lower throughout (generally <95% UFBOOT although 98% at 50% sites remaining) and dropped precipitously when 30% of the sites remained.

Random subsampling of 50% of the genes in 5 jackknife replicates maintained robust support for *Meteora*+Hemimastigophora but not for *Ancoracysta+Meteora*+Hemimastigophora (Fig S4H). The gene concordance factor (gCF^19^) for *Meteora*+Hemimastigophora (8.65%) was similar, or higher, than that of several accepted supergroups like Sar (9%), CRuMs (4.35%) and Amorphea (1.2%). By contrast, *Ancoracysta*+*Meteora*+Hemimastigophora was recovered in <1% of the single gene trees (gCF 0.87%; Fig S4F). We infer that the phylogenetic signal for the *Meteora*+Hemimastigophora relationship is broadly distributed across genes.

Overall, the *Meteora*+Hemimastigophora association remained robust through tests for biases from subsets of genes and sites, and, notably, those for compositional bias (i.e. the recoding analyses). On the other hand, *Ancoracysta*+*Meteora*+Hemimastigophora was poorly supported in these tests, especially for compositional bias. The position of Provora, represented here by *Ancoracysta,* remains unresolved by our study, as in prior examinations ^4,20^.

### The Meteora mitochondrial genome is gene-rich

We completely sequenced the mitochondrial genome of *Meteora sporadica* LBC3 (*Met* mt-genome). The genome is a circular mapping molecule of 94.9 kbp (94,877 bp) with a G+C content of 28.8%. The *Met* mt-genome encodes a total of 79 genes (38 duplicated) including 50 protein-coding genes, 2 functionally unidentified open reading frames (ORFs), and 27 RNA genes (*rnl*, *rns*, *rrn5* and 24 tRNAs) (Fig 4A). The tRNAs recognize 44 codons that together code for all 20 amino acids, but no stop-codon-recognizing tRNAs were found. No recognizable mobile elements, introns, or split-genes were detected in the genome. The genome contains a pair of inverted repeats of 33.0 kbp (32,998 bp) that are separated by unique regions of 2,143 bp and 26,739 bp. The shorter unique region encodes *ccmF* bounded by two tRNAs, while the larger one has 25 protein-coding genes and 14 tRNAs.

**Figure 4.**
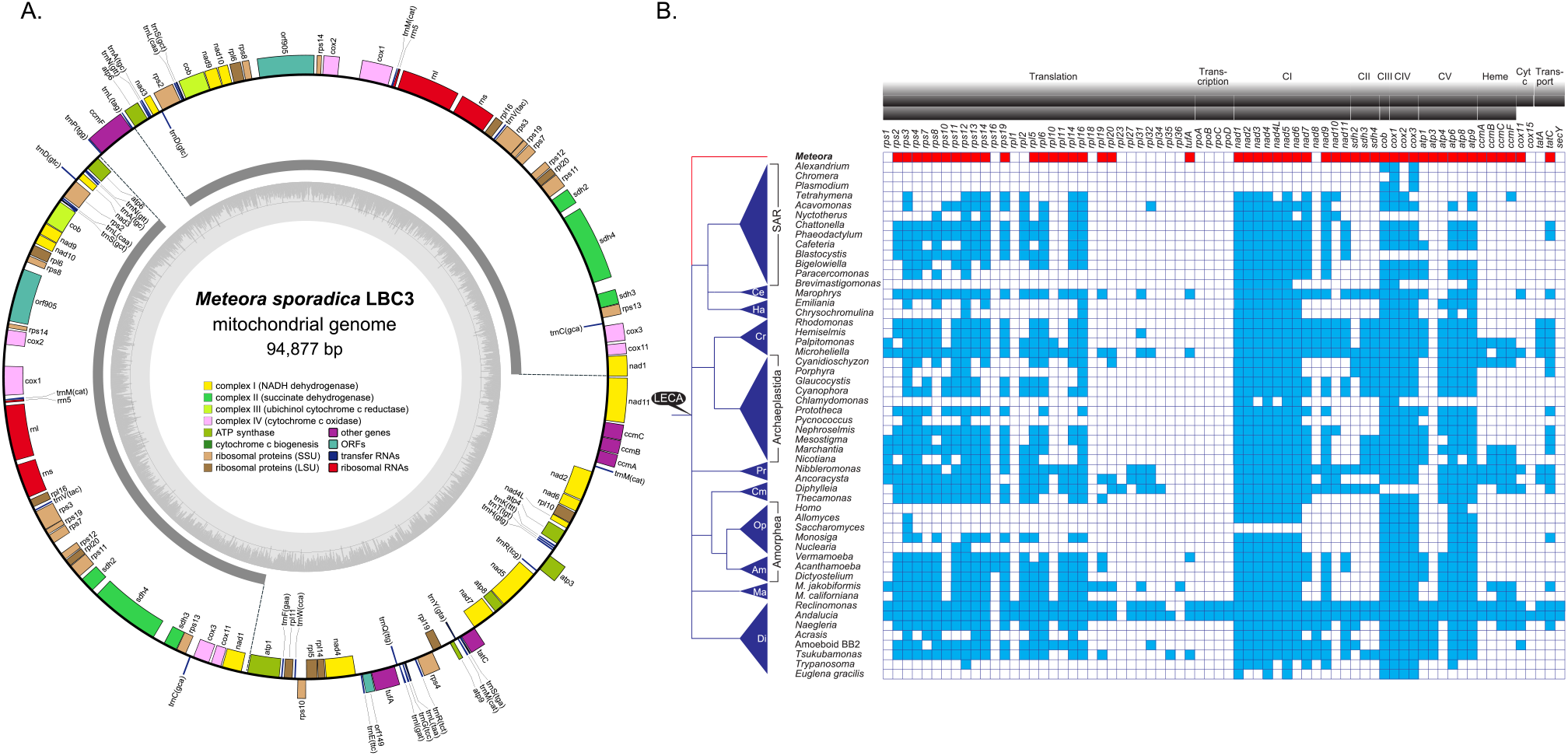
A gene-rich mitochondrial genome in *Meteora*. A) The map of the *Meteora sporadica* LBC3 mitochondrial genome with genes color-coded as to their function and the GC% indicated in grey. Genes on the outside of circle are transcribed in the clockwise direction. The inverted repeats are indicated in dark grey segments. B) Protein-coding capacity of mitochondrial genomes across eukaryotes. Presence and absence of corresponding genes from selected eukaryotes are shown by blue or white boxes respectively. The presence and absence of genes are derived from ^24^ and updated with additional lineages. Phylogenetic relationships between eukaryotes are based on but showing the eukaryotic root as a polytomy ^16,25,84^. The *Meteora* branch is indicated in red. Abbreviations: Al - Alveolata, St - Stramenopila, Rh - Rhizaria, Ce - Centrohelea, Ha - Haptophyta, Cr - Pancryptista, Re – Rhodophyta (‘Reds’), Gl - Glaucophyta, Ch - Chlorophyta, Pr - Provora, Cm - CRuMs, Op - Opisthokonta, Am - Amoebozoa, Ma - Malawimonadida, Di - Discoba. CI-CV - electron transport chain complex I–V.

The *Met* mt-genome is among the most gene-rich yet documented, with only jakobids (60-66 genes), *Microheliella maris* (53 genes), nibblerids (51 genes), and *Diphylleia rotans* (51 genes) encoding larger gene sets ^16,21–24^. The 24 unique genes for the respiratory chain complexes in the *Met* mt-genome is the most complete mt-encoded respiratory complex gene set, together with *Microheliella*, outside of jakobids ^21,22,25^(Fig 4B). This includes *atp3*, the first such gene recorded outside of Discoba ^21,22,26–30^.

Other rare genes in the *Met* mt-genome include *tufA* and *cox11*. These have previously only been seen together in the mitochondrial genomes of some Discoba (*Tsukubamonas globosa* and jakobids), the centrohelid *Marophrys* sp., and *Microheliella maris*, a protist which branches basal to Cryptista ^25,31,32^. The mitochondrial genomes of *Diphylleia rotans,* a member of CRuMs, and members of the provoran taxon *Nibbleromonas* encode only *cox11* ^16,24^, while the amoebozoan *Vermamoeba vermiformis* mt-genome encodes only *tufA* (GenBank accession number: *<u>GU828005</u>;* unpublished).

We detected a full set of type I cytochrome *c* maturase genes (*ccmABCF*), which, in many eukaryotes, has been functionally replaced by the eukaryote-specific nuclear-encoded type III system, holocytochrome *c* synthase (HCCS) ^33^. System I is presumed to be ancestral to all eukaryotes and has been replaced over time by HCCS, but it is unclear whether this happened as a single event or multiple times ^31,33,34^.

### Conclusions

Phylogenomic analyses convincingly show *Meteora* as a sister group to Hemimastigophora. This seems remarkable based on their morphology and basic life history. As shown here, *Meteora* cells are completely aflagellate bacterivores, whereas hemimastigotes are multi-flagellated cells that prey on microbial eukaryotes ^2,35^. Both exhibit symmetry, which is relatively uncommon among unicellular eukaryotes; however, hemimastigotes have diagonal symmetry and are essentially a constant shape, whereas *Meteora* is predominantly bilaterally symmetrical and is highly plastic, breaking and re-establishing symmetry in the arrangement of the arms. While these groups seem to have little in common, established eukaryotic supergroups like Sar and CRuMs also encompass a bewildering variety of morphologies and lifestyles. Sar encompasses fungal-like, flagellated, and amoeboid forms, and even large macroscopic algae^36^. Although CRuMs is represented by fewer than 10 described species, these range from small filose amoebae to bacterivorous nanoflagellates to larger eukaryovorous flagellates^37^.

This finding resonates with the high rate of discovery of novel eukaryotes, and indeed entire new phylum- and supergroup-level lineages, over recent years^1,2,4,15,20^. As with all other recently discovered major lineages^1^, *Meteora* is a free-living heterotrophic protist, underlining the importance of pursuing this category of organism in efforts to catalogue deeper eukaryote diversity. It also illustrates that the first known representative of a major clade (i.e. hemimastigotes in *Meteora*+Hemimastigophora) need not reflect the morphology or biology of the rest of the group. In particular, environmental lineages (i.e. groups known only from molecular data) may not necessarily be similar to their morphologically-characterised relatives. Both the remarkable cellular architecture and unexpected phylogenetic placement of *Meteora sporadica* suggest that the staggering diversity of microbial eukaryotes is far from fully understood, and will continue to surprise us.

## Extended METHODS

### Isolation and cultivation

Samples were obtained from subtidal/intertidal sediments. For SRT610, a sample from Nikadori fishing port, Okinawa, Japan (24° 49’ 10.52’’, 125° 16’ 47.77’’) was enriched in ESM medium at 20°C. An individual cell was picked with a drawn-out glass micropipette and placed in ESM medium. For LBC3, a sample from Playa la Boca, Cuba (21° 35’ 24.99’’, -77° 5’ 35.33’’; kindly provided by Claire Burnard), was enriched in seawater + LB medium at room temperature (21°C). An individual cell on a flake of biofilm was picked by micropipette and placed in 0.1% LB in autoclaved natural seawater medium. Both isolates were subsequently maintained in tissue culture flasks with unidentified co-cultured bacteria at 20°C (SRT610) or 16°C (LBC3), and transferred every 2 weeks.

### Light microscopy

For SRT610, aliquots of culture were mounted on slides with coverslips and imaged on Zeiss Axio imager A2 microscope (Carl Zeiss AG) with an Olympus DP74 CCD camera (Olympus), while aliquots of LBC3 culture were incubated on sealed slide preparations overnight and imaged on Zeiss AxioVert 200M with an AxioCam ICc5 camera (Carl Zeiss AG). Downstream image processing and analysis was done in FIJI^38,39^.

### Transmission electron microscopy

Culture flasks of isolate SRT610 were scraped and removed cells were collected by centrifugation (1000 *g*, 15 min). The concentrated material was mounted on copper grids and plunged rapidly into liquid propane. The frozen pellets were then plunged into liquid nitrogen for several seconds, then placed in acetone with 2% osmium tetroxide at −85 °C for 48 h. The fixing solution was then kept at −20 °C for 2 h and at −4 °C for 2 h. The pellets were rinsed with acetone three times, and were then embedded in agar low viscosity resin R1078 (Agar Scientific Ltd, Stansted, England). The resin was polymerized at 60 °C for 12 h. Ultrathin sections were prepared on a Reichert Ultracut S ultramicrotome (Leica, Vienna, Austria), double stained with 2% uranyl acetate and lead citrate, and observed using a Hitachi H-7650 electron microscope (Hitachi High-Technologies Corp.) equipped with a Veleta TEM CCD camera (Olympus).

### SSU rDNA phylogenetics

DNA was extracted from SRT610 using the DNeasy Plant Mini kit (Qiagen), and from LBC3 using the DNeasy Blood & Tissue kit (Qiagen). The SSU rDNA of SRT610 was amplified by PCR using forward primer 18F (5′-AAC CTG GTT GAT CCT GCC AG-3′) and reverse primer 18R (5′-CYG CAG GTT CAC CTA CGG AA-3′) at 55°C annealing temperature for 35 cycles. The SSU rDNA of LBC3 was obtained by semi-nested PCR, with initial amplification using forward primer EukA (5′-AACCTGGTTGATCCTGCCAGT-3′) and reverse primer 1498R (5′-CACCTACGGAAACCTTGTTA-3′) at 63°C annealing temperature for 35 cycles, followed by secondary amplification with forward primer 82F (5′-GAAACTGCGAATGGCTC`-`3′) and reverse primer 1498R at 63°C for 25 cycles. The sequences were obtained by Sanger sequencing, with some PCR product from LBC3 being gel-extracted prior to sequencing (QIAquick Gel Extraction kit; Qiagen).

The *Meteora* sequences were added to a global eukaryotic SSU alignment (derived from the reference SSU dataset for the environmental analysis in Lax et al. 2018^2^) via profile alignment in SeaView^40,41^. The alignment was further augmented for taxon sampling with additional environmental sequences from NCBI and Jamy et al. (2020)^42^, corrected manually, then masked via gblocks^43^ followed by manual correction to yield a 1187 site alignment across 173 taxa. This was subject to phylogenetic analyses in RAxML^44^ (raxmlHPC-PTHREADS-SSE3 v. 8.2.6) under the GTR+Γ+I model with 50 starting trees and 1000 non-parametric bootstraps.

### Combined SSU and LSU rDNA phylogenetics

Source alignments for SSU and LSU rDNA from Jamy et al. (2020)^42^ were expanded for broader taxon selection using publicly available data in NCBI nt, or extracted from published transcriptome and genome assemblies using barrnap^45^ v. 0.9. *Meteora* LBC3 LSU rDNA was extracted from the transcriptome using barrnap and concatenated with the SSU rDNA mentioned above. Site selection was performed on each alignment using g-blocks ^43^ in SeaView^40^ followed by manual curation, then the SSU and LSU rDNA alignments were concatenated for a total of 3051 sites. The phylogeny was inferred via RAxML^44^ (raxmlHPC-PTHREADS-SSE3 v. 8.2.6) under the GTR+Γ+I model with 50 starting trees and 1000 non-parametric bootstraps.

### Environmental sequence analysis

We searched 14 153 628 publicly available V4 and V9 sequences from TARA Oceans (V9)^46^, VAMPS (V9)^47^, MetaPR2 (“Biomarks”) (V4)^48,49^, deep sea sediments (V9)^50^, Malaspina (V4)^51^, neotropical^14^ and temperate^52^ soil metatranscriptomes (V4), and the Cariaco basin oxic-anoxic gradient (V4)^53^ for sequences very similar to *Meteora*. The V4 and V9 regions of the *Meteora* LBC3 SSU rDNA were extracted and used to query the respective databases using BLASTn with a sequence identity threshold of 80%. The collected sequences were then aligned using PaPaRa^54^ against a 1187 site, 173 taxon reference SSU rDNA alignment derived from Lax et al. (2018)^2^, manually curated through MUSCLE^41^ profile alignments in SeaView^40^, and augmented for taxon sampling with additional environmental sequences from NCBI and Jamy et al. (2020)^42^ (Fig S3A). Phylogenetic placements were inferred via RAxML-EPA^55^. Output was analysed in R using ggtree^56^ and filtered with a likelihood-weight ratio threshold of 0.5.

### Transcriptome assembly

For RNA extraction, SRT610 cells grown in culture flasks were dislodged by scraping and collected by centrifugation at 3000 *g* for 10 minutes at room temperature. Total RNA was extracted using TRIzol (ThermoFisher) following the manufacturer’s instructions. The cDNA library construction and paired-end sequencing (125 bp per read) with Illumina HiSeq2500 were performed at Eurofins Genomics (Tokyo, Japan). Read quality was inspected using FastQC^57^, adaptors clipped and reads trimmed with Trimmomatic v.0.30^58^ (LEADING:3 TRAILING:3 SLIDINGWINDOW:4:15 CROP:160 MINLEN:36), and assembly performed in Trinity^59^ v2.2.0.

For LBC3, RNA was extracted from culture grown in 0.1% LB in sterile seawater on Petri plates (15 cm diameter), scraped and spun 30 min at 2500 *g* and 16°C, followed by adding 15 mL TRIzol (ThermoFisher) to 5 mL of resuspended pellet. Then, 3 mL of chloroform was added and phase separation obtained by centrifugation for 30 min at 4500 *g* at 4°C. The aqueous phase was removed and further treated as per manufacturer’s instructions. The RNA was further purified with a phenol:chloroform extraction and treated with DNase. Quantity was assessed by Qubit (ThermoFisher). The sequencing library was prepared using the NEBNext Poly(A) mRNA Magnetic Isolation Module (NEB #E7490; New England Biolabs), and sequenced on Illumina MiSeq with 2 x 250(V2 kit) bp reads, indexed with Illumina adaptors i703 and i503 (multiplexed with an undescribed metamonad with adaptors i704 and i504). Read quality was inspected using FastQC^57^, adaptors clipped and reads trimmed with Trimmomatic^58^ v.0.30 (LEADING:3 TRAILING:3 SLIDINGWINDOW:4:15 CROP:160 MINLEN:36) and assembled with Trinity^59^ v.2.0.2. To remove most cross-contamination from multiplexed samples, we used a custom script (M. Kolisko, Institute of Parasitology Biology Centre, Czech Academy of Sciences, České Budějovice) and then reassembled in Trinity.

Transcriptome completeness was assessed by BUSCO^60^ v3.0.2 using eukaryote_odb10 dataset. This yielded 231/255 complete BUSCOs (13 fragmented) for SRT610 and 211/255 complete BUSCOs (24 fragmented) for LBC3. Of the 254 phylogenomic marker genes (see below), 236 from each isolate were present in the final alignment, with 90.2% and 88.5% site occupancy for SRT610 and LBC3, respectively.

### Phylogenomic dataset assembly

The 351 gene phylogenomic dataset from Lax et al. (2018)^2^ (based originally on Brown et al. 2018^37^) was expanded by adding the two *Meteora* isolates, plus selected subsequently sequenced taxa including *Ancoracysta*^20^, telonemids^15^, and the three *Rhodelphis* transcriptomes^4^ via a custom pipeline^37^. Telonemids, *Rhodelphis*, and *Meteora* were added (and Hemimastigophora re-added) using a custom script that enables multiple candidate genes per transcriptome to be selected and added at once, up to 4 in this case. After addition, each gene was re-aligned with MAFFT-linsi^61^, trimmed with BMGE^62^ (-h 0.5, -g 0.2, -m BLOSUM30), and phylogenies inferred under the LG4X+Γ model^63^ in IQ-TREE v1.5.5^64^, then manually inspected for paralogues, contaminants, lateral gene transfers, and signs of deep paralogies within the base dataset. Sequences marked for deletion were removed using a custom script. Where deep paralogies were detected that affected the whole gene tree, we discarded the gene from the dataset, resulting in a final phylogenomic dataset of 254 genes (listing on Datadryad). The single gene alignments were filtered using PREQUAL^65^ with -filterthresh 0.95 (0.28% masked), then trimmed with BMGE (-h 0.5, -g 0.2, -m BLOSUM30) and concatenated, for a final alignment of 70471 sites. The taxa were subsampled to produce a 108-taxon dataset aiming to broadly represent eukaryote diversity, and a 66-taxon dataset for computationally intensive analyses (see listing on Datadryad). In both cases, phylogenetically redundant taxa were removed, with retention of higher coverage and more slowly evolving taxa where possible.

### Phylogenomic analyses

An initial phylogeny was inferred from the concatenated 254-gene, 108-taxon dataset in IQ-TREE v1.5.5^64^ using the LG+C20+F+Γ model, with support assessed via UFBOOT bootstrap approximation (1000 replicates) in IQ-TREE^66^. Next, a phylogeny was inferred from the subsampled 66-taxon dataset under the LG+C60+F+Γ model, then used as a guide tree for the 60 custom profile site-heterogeneous mixture model LG+MAM60+Γ^67^ (hereafter referred to as “MAM60”) using the program MAMMaL^68^, with support values generated via UFBOOT bootstrap approximation in IQ-TREE^66^. MAM60 was preferred over C60 by AIC (7324035 - 7302542 = 21493) and BIC (7325959 – 7314353 = 11606). A site-heterogeneous mixture model approximation method, PMSF^69^, was used to generate 200 non-parametric bootstrap trees using the MAM60 tree as the guide tree. A Bayesian phylogeny was inferred using the CAT+GTR model in Phylobayes^70^ v. 1.8 via 4 chains, with 1.1 x 10^4^ cycles and a burn-in of 500. Three chains converged, but the unconverged chain (1) was identical in all respects directly relevant to the placement of *Meteora* (compare Fig S4I and Fig S4J).

The Hemimastigophora+*Meteora* relationship was interrogated further via downstream analyses based on the 66-taxon dataset. A step-wise removal of fastest evolving sites (Fast Site Removal – FSR) was done in 10% increments using Phylofisher v.0.1.20^71^ and corresponding phylogenies inferred under MAM60 with UFBOOT support. Support values for relationships of interest were summarised via a custom script. A ‘no long-branching taxa’ (nLB) alignment was produced by determining the outlier long branches via a custom script (L. Eme; CNRS at Université Paris-Sud, France), in this case *Tetrahymena*, *Diplonema*, and *Bodo*. This dataset, along with one with *Ancoracysta* removed (noAnco) was used to infer a phylogeny under the MAM60 model.

To test whether the Hemimastigophora+*Meteora* relationship was the result of a few outlier genes, two analyses were conducted: gene jack-knifing and gene concordance factor (gCF^19^) calculation. 5 gene-jack-knifing replicate alignments of 50% of the genes (following recommendations for adequate statistical power in Brown et al. 2018^37^) were generated using random_sample_iteration.py utility in Phylofisher^71^, and corresponding phylogenies inferred under MAM60 in IQ-TREE with statistical support from 1000 UFBOOT replicates. Single gene trees were estimated under MAM60 in IQ-TREE v1.5.5 for each of the 254 individual gene alignments and gCF calculated in IQ-TREE v2.0^72^.

To test for biases arising from sequence composition, two recoding approaches were used. The Susko and Roger set of 4 amino acid classes (SR4^17^) was used to reduce the amino acid alphabet. Additionally, a set of 4 amino acid classes that minimises compositional differences between sequences was determined via minmax-chisq^18^. In both cases, these schemes were used to recode the amino acid alignment as well as the 60 category MAMMaL model definition via custom scripts (see DataDryad), and then a phylogeny was inferred under the GTR+[4binCustomModel]+R6 model in IQ-TREE 2.0, with support values inferred from 1000 UFBOOT replicates. Trees were formatted using the Ete3 toolkit^73^.

### DNA extraction for mitochondrial genome sequencing

Isolate LBC3 was grown in K media^74^ with 0.3% LB at room temperature for 3-4 days until most bacteria were consumed and the culture dish was dense with cells. *Meteora* cells from two litres of culture (50 150 mm x 15 mm Petri dishes) were harvested by careful decanting of 90% of the volume. The cells were collected by scraping and pooled in 50 mL Falcon tubes, then pelleted by centrifugation in a swing-out rotor at 2000 *g*, 10 min, 20°C. The pelleted cells were resuspended in artificial sterile seawater (ASW) and re-pelleted by centrifugation as above in a 15 mL Falcon tube. The cell pellet was resuspended in 4 mL of ASW and 1 mL aliquots were pelleted for 2 min at 16000 *g*, 4°C. The dry pellets were frozen at -80°C or used directly for long-read DNA extraction.

Cells for short-read sequencing were harvested from two 175 cm^2^ culture flasks grown for 3 weeks at room temperature. The cultures were harvested by first carefully decanting 90% of the volume and then dislodged using a cell scraper. The cells were pelleted as described in the previous paragraph.

DNA for long-read and short-read sequencing was purified using the MagAttract HMW gDNA kit (Qiagen) using the tissue lysis protocol. DNA for long-read sequencing was additionally purified on the GenomicTip G/20 column (Qiagen) by the manufacturer’s protocol. Sample quality and quantity were assessed by agarose gel electrophoresis, a nano spectrophotometer and the Qubit™ dsDNA BR Assay Kit (Thermo Fisher Scientific).

### Short-read DNA sequencing

DNA for Illumina short-read sequencing was submitted to Génome Québec for shotgun-library construction using the Illumina TruSeq LT kit. The libraries were sequenced on an Illumina HiSeq X using 150 bp paired reads. Illumina reads were quality checked using FastQC v.0.11.5 (http://www.bioinformatics.babraham.ac.uk/projects/fastqc) and trimmed using Trimmomatic v0.36^75^. Short-reads derived from the mitochondrial genome were recruited by mapping reads using Bowtie2^76^ against a circular mapping mitochondrial genome contig assembled using default settings in Abruijn v1.0^77^.

### Long-read DNA sequencing

Long-read data was generated in two sequencing runs. Oxford Nanopore libraries were prepared with ligation sequencing kits SQK-LSK109 and SQK-LSK110 (sequencing runs 1 and 2, respectively) and sequenced on MIN106D flow cells (R9.4.1). Base-calling was performed using Guppy v5.0.11 using the SUP (super high accuracy) model. Adapters and chimeric reads were removed using Porechop v0.2.4 (default settings with *--discard_middle* option).

Long reads were assembled with Flye v2.9 in metagenome mode (*-meta* flag)^78,79^. The contig containing the mitochondrial genome was identified by mapping previously identified mitochondrial short reads to the long-read assembly with HISAT2 v2.2.1^80^. Repetitive regions in this contig were collapsed – to generate a single circular contig with resolved repeats, NGMLR v0.2.7 ^81^ was used to identify the long reads that mapped only to the mitochondrial contig, and a new assembly was generated using those reads with Flye v2.9 under default settings.

The assembly was polished using two rounds of long-read polishing with Medaka v1.7.2 followed by one round of short-read polishing with Pilon v1.24 ^82^. Short reads were then mapped to the polished assembly (HISAT2 v2.2.1) and the few remaining sequencing errors were identified using a genome browser (Tablet v1.21.02.08) ^83^ and manually corrected.

MFannot v1.36 (https://megasun.bch.umontreal.ca/apps/mfannot/) was used for gene prediction and annotation using the standard genetic code. Annotations and gene boundaries were inspected in Tablet, and any missing annotations were added manually.

## Acknowledgements

We would like to acknowledge Claire Burnard for providing the sample for isolate LBC3, and André Comeau (IMR, Dalhousie U.) for help with Illumina sequencing. This study was supported by the Japan Society for the Promotion of Science (JSPS) KAKENHI Grant Numbers: 13J00587, 18J02091, by NSERC discovery grant 298366-2019 (to AGBS), and NSERC Discovery grant RGPIN-2022-05430 (to AJR)

## Author Contributions

Conceptualization, YE, ST, KI and AS; Investigation, YE, ST, KW and JJ-H; Formal Analysis, YE, ST, KW and JJ-H; Visualization, YE, ST, KW and JJ-H; Supervision, AR, KI and AS; Funding Acquisition, AR, KI and AS; Writing – Original Draft, YE, KW, JJ-H and AS; Writing – Review & Editing, all authors.

## Data availability

All data will become available in GenBank once the manuscript has been accepted for publication by a peer-reviewed journal.

Transcriptome assemblies, alignments for rDNA analyses, and alignments for phylogenomic analyses, plus all videos of live *Meteora sporadica*, will become available on Datadryad once this manuscript is accepted for publication in a peer reviewed journal.

## Supplementary video descriptions

**Supplementary Video S1.** Real time video of *Meteora sporadica* isolate LBC3 gliding, followed by a video of one cell gliding into another and the long axis projection bending. Note motion of the ‘arms’.

**Supplementary Video S2.** Real time video of a specimen of *Meteora sporadica* isolate LBC3 with a more complex arrangement of projections.

**Supplementary Video S3.** Real time video of an ‘armless’ specimen of *Meteora sporadica* isolate LBC3 gliding, showing that the gliding motility does not depend on the ‘arm’ motion.

**Supplementary Video S4.** Real time video of a *Meteora sporadica* isolate LBC3 cell picking up a food bacterium; 2x sped up video of several cells feeding; real time video of a cell, already carrying a prey bacterium, firing an extrusome at another bacterium (unsuccessfully).

---

**Figure S1.**
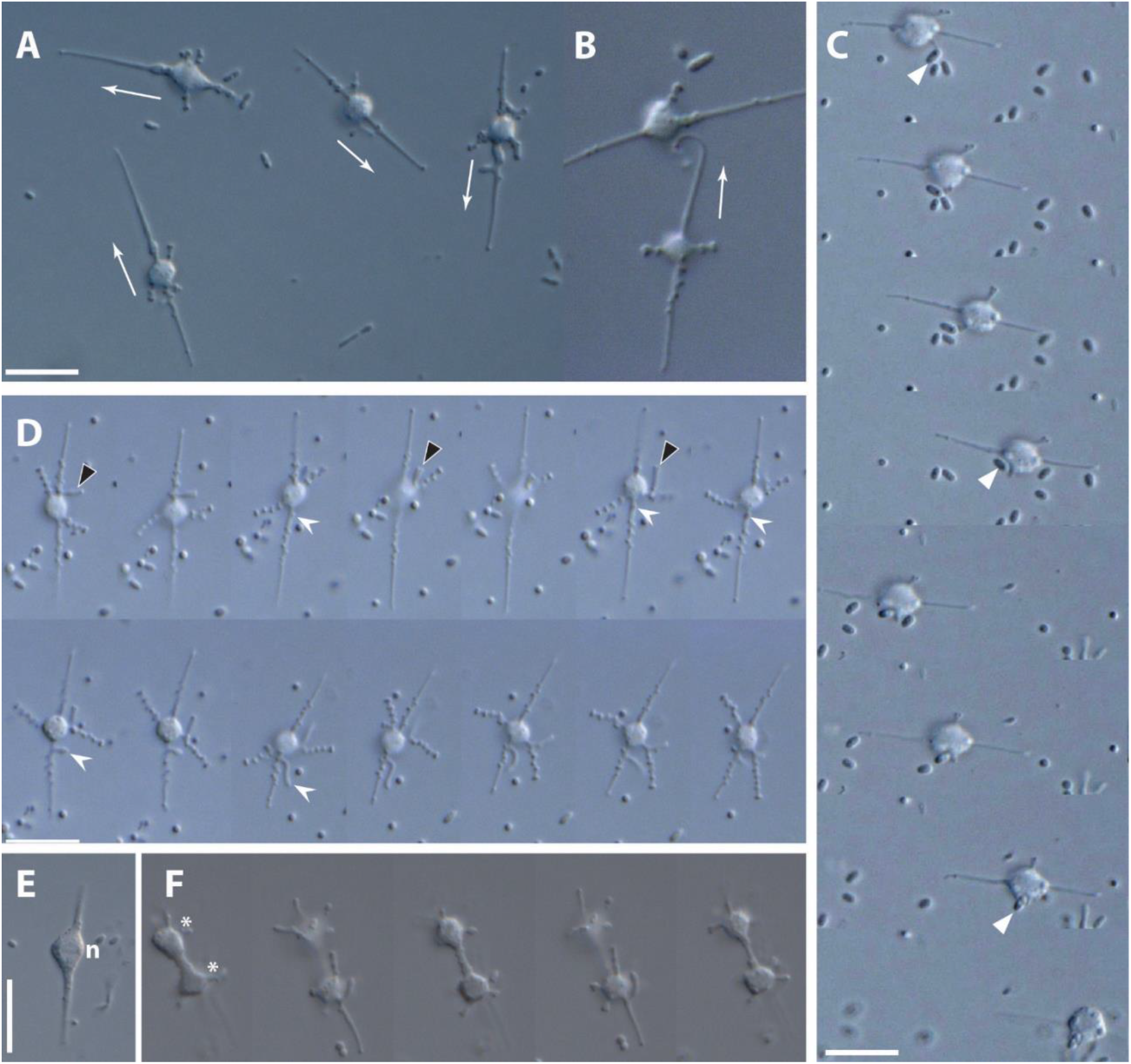
Additional light micrographs of *Meteora sporadica* isolate LBC3. A) General view of four cells of *Meteora sporadica* isolate LBC3, showing their direction of movement (arrows), and variety in lateral ‘arm’ morphology. B) Collision between two cells (arrow shows direction of movement) showing the bending of the long axis. C) Feeding on a bacterium. A granule on an arm plays a role in contacting and attaching the bacterium (arrowhead), which is then moved towards the cell body proper and phagocytosed there. D) Series showing the behaviour of cytoskeletal elements and surface granules. Protrusions can jump between the long axis and the arms across the surface of the cell body proper (black arrowhead). Regions of the axes associated with surface granules can protrude outwards, sometimes rapidly (white barbed arrow), and later fuse with the long axis (not shown). E) Cell in early division, arms retracted, as the cell body proper, containing the nucleus (n) in mitosis, gradually moves up and down along the long axis. F) Later stage of another dividing cell. Cells separate along the long axis and gradually begin to reconstitute arms starting from this stage. Scalebars: 10 µm. Videos corresponding to B and C are in supplementary Videos 1 and 4, respectively. Videos for A, E and F are available on DataDryad.

**Figure S2.**
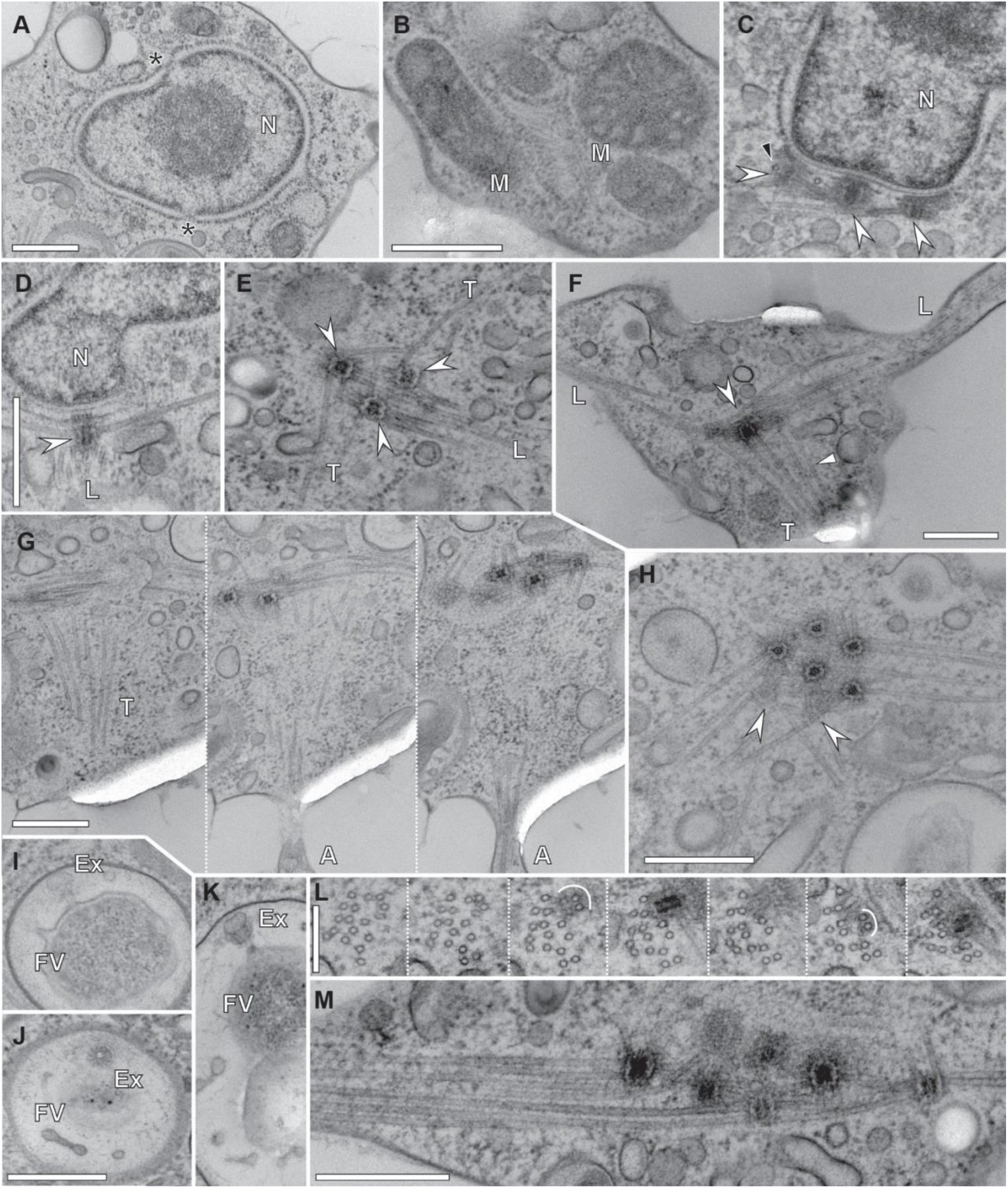
Additional transmission electron micrographs showing ultrastructure of *Meteora sporadica* isolate SRT610. A) Nucleus (N) with two pores visible (asterisks). B) Mitochondrial (M) section showing cristae in longitudinal (left) and transverse (right) sections. C) MTOCs (barbed arrowhead) associated with the nucleus (N), with cross sections of perpendicularly-oriented microtubules emerging nearby (black arrowhead). D) Another section through an MTOC (barbed arrowhead) detailing its association with the nuclear (N) envelope, as well as an oblique section through the longitudinal microtubular bundle (L). E) A small cluster of MTOCs (barbed arrowheads) with the emerging longitudinal bundle (L) and transverse (T) microtubules. F) General view of the cell body with the longitudinal bundle (L) as well as transverse microtubules (T, arrowhead) emerging from the cluster of MTOCs (barbed arrowhead). Adjacent section in series containing Fig 2B. G) Series (70 nm steps) following transverse microtubules (T) from the cluster of MTOCs to the start of the lateral arm (A). H) A cluster of 7 MTOCs, two of them in grazing section (barbed arrowheads). I-K) Detail of discharged extrusomes (Ex) inside food vacuoles (FV) containing bacteria. The extrusome in J is in cross section. L) series following the longitudinal bundle of microtubules through two MTOCs of a cluster. Populations of microtubules emerging in the adjacent section are indicated by a white arc. Connective material can be seen between some microtubules in the bundle. M) Longitudinal section of the longitudinal bundle and MTOCs in a different cell, to the same scale as L, as a reference. Scale bars: A, B-C (in B), D-E (in D), F, G, H, I-K (in J) all 500 nm, L) 250 nm, M) 500 nm.

**Figure S3.**
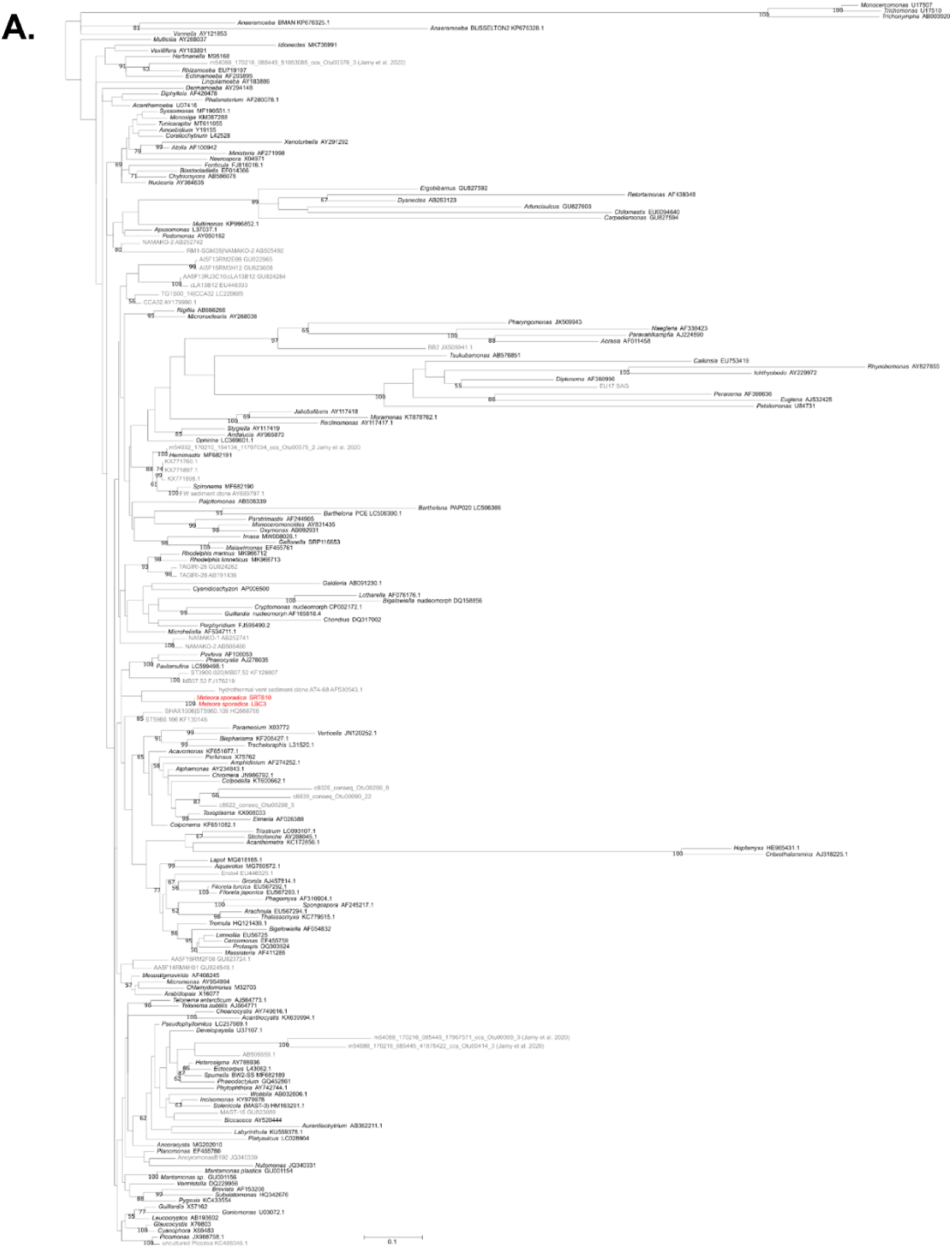
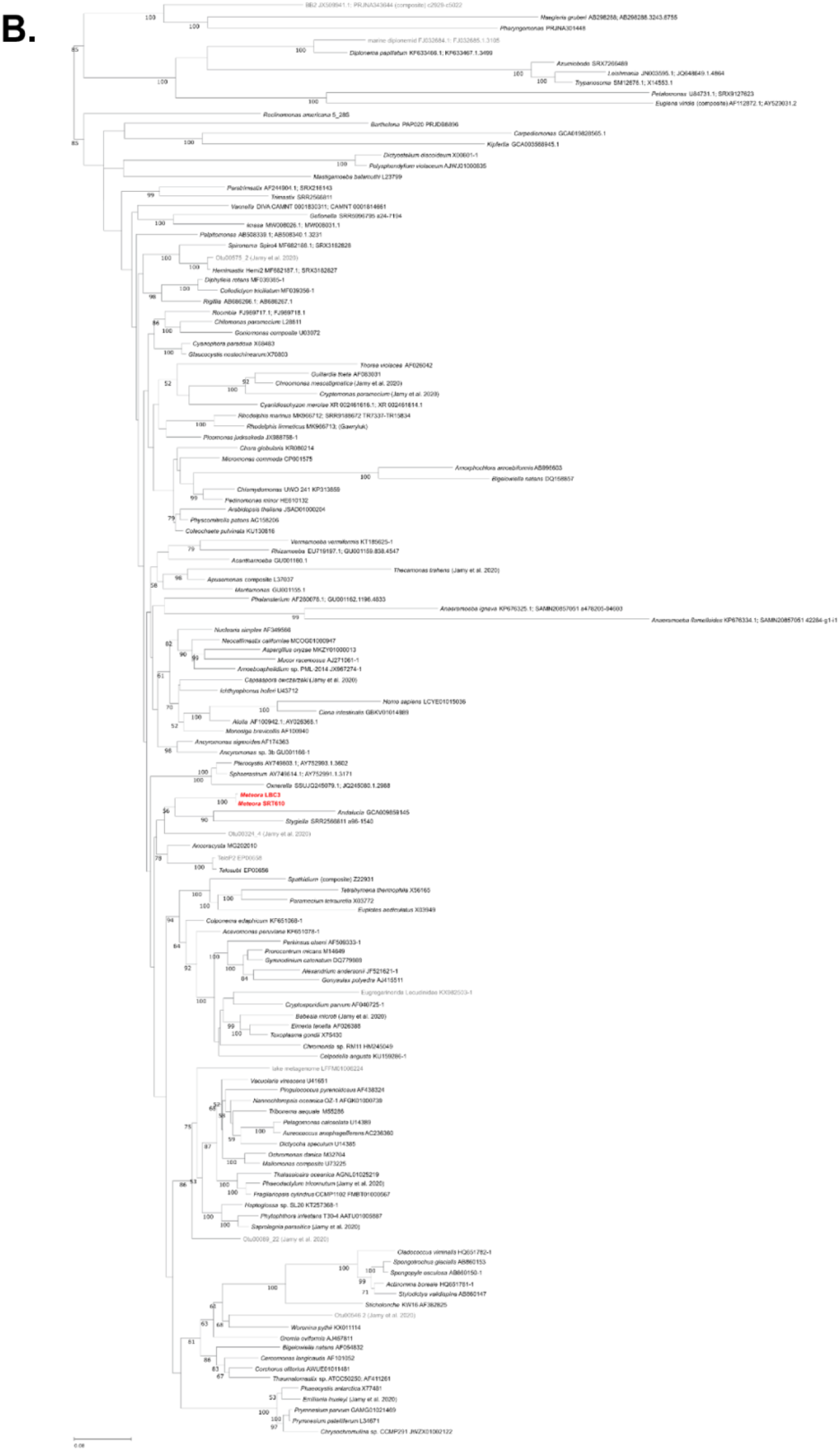
Position of *Meteora sporadica* in rDNA phylogenies. A) SSU rDNA phylogeny representing eukaryote-wide diversity, for use as the reference tree for environmental sequence placement analyses. Alignment contains 1187 sites across 173 taxa. Tree inferred under the GTR+Γ+I model with 1000 non-parametric bootstrap replicates. *Meteora* sequences highlighted in red. B) SSU-LSU rDNA phylogeny inferred from 3051 sites in final concatenated alignment, across 137 taxa, under the GTR+Γ+I model with 1000 non-parametric bootstrap replicates. *Meteora* sequences are highlighted in red.

**Figure S4.**
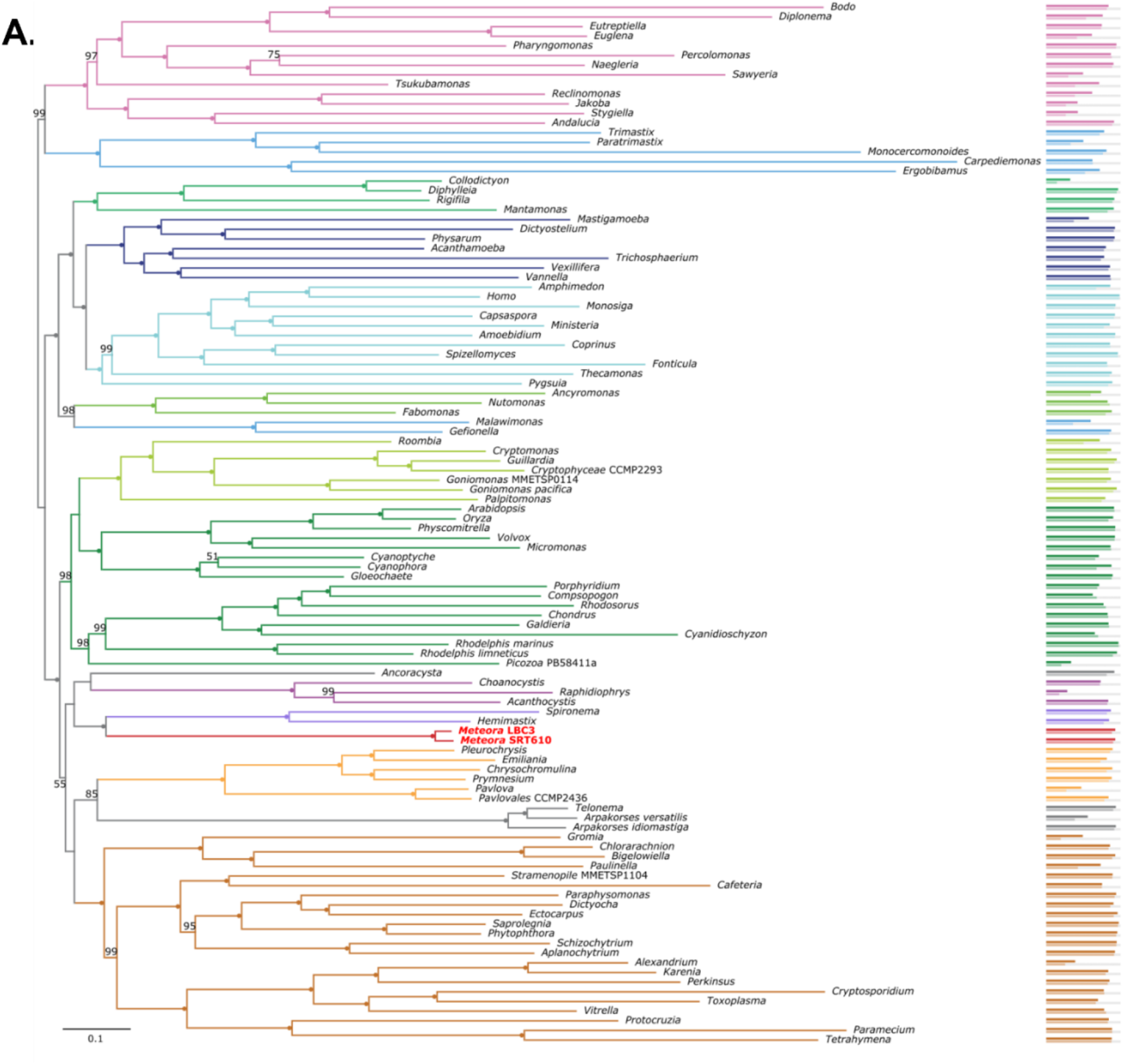
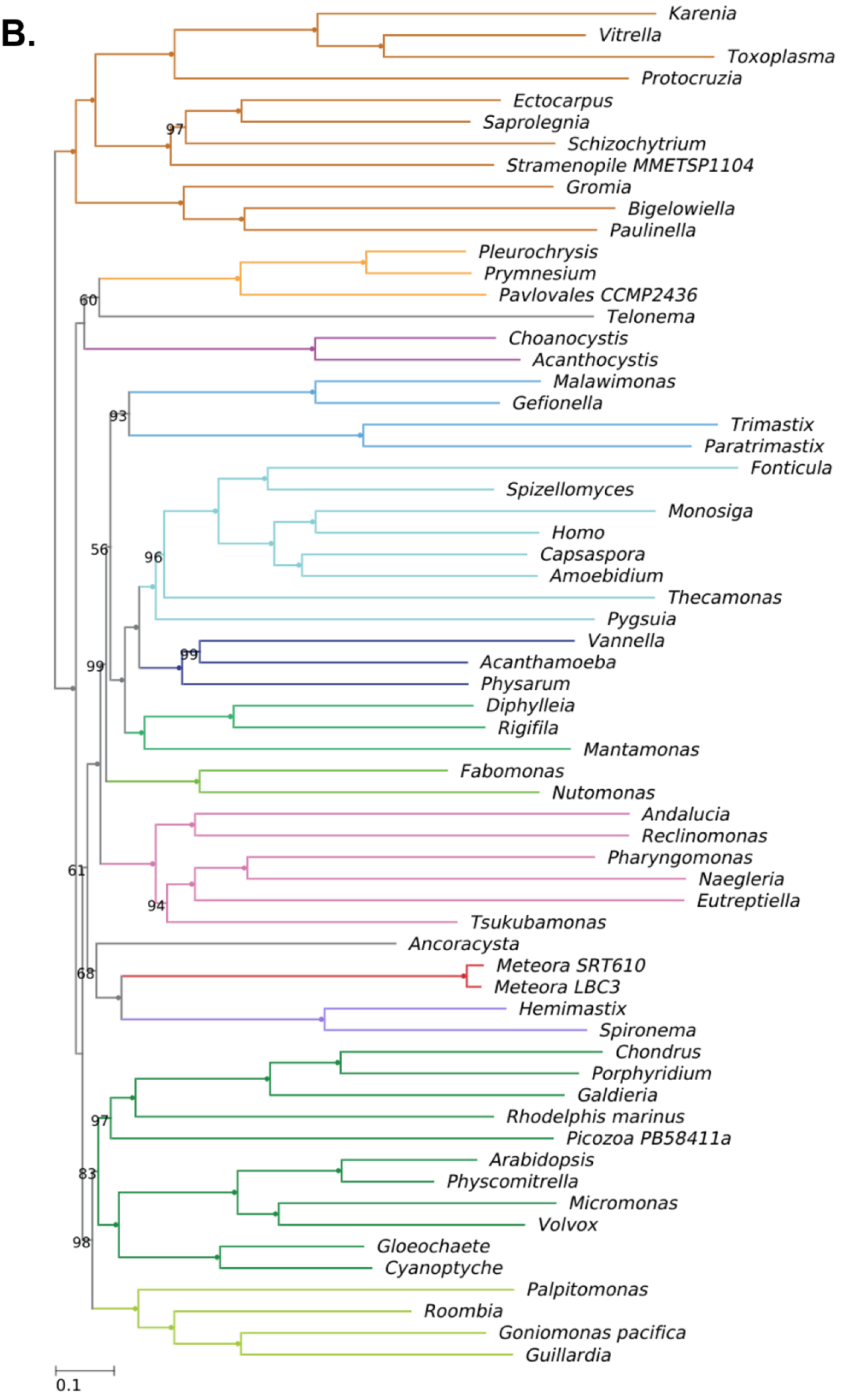
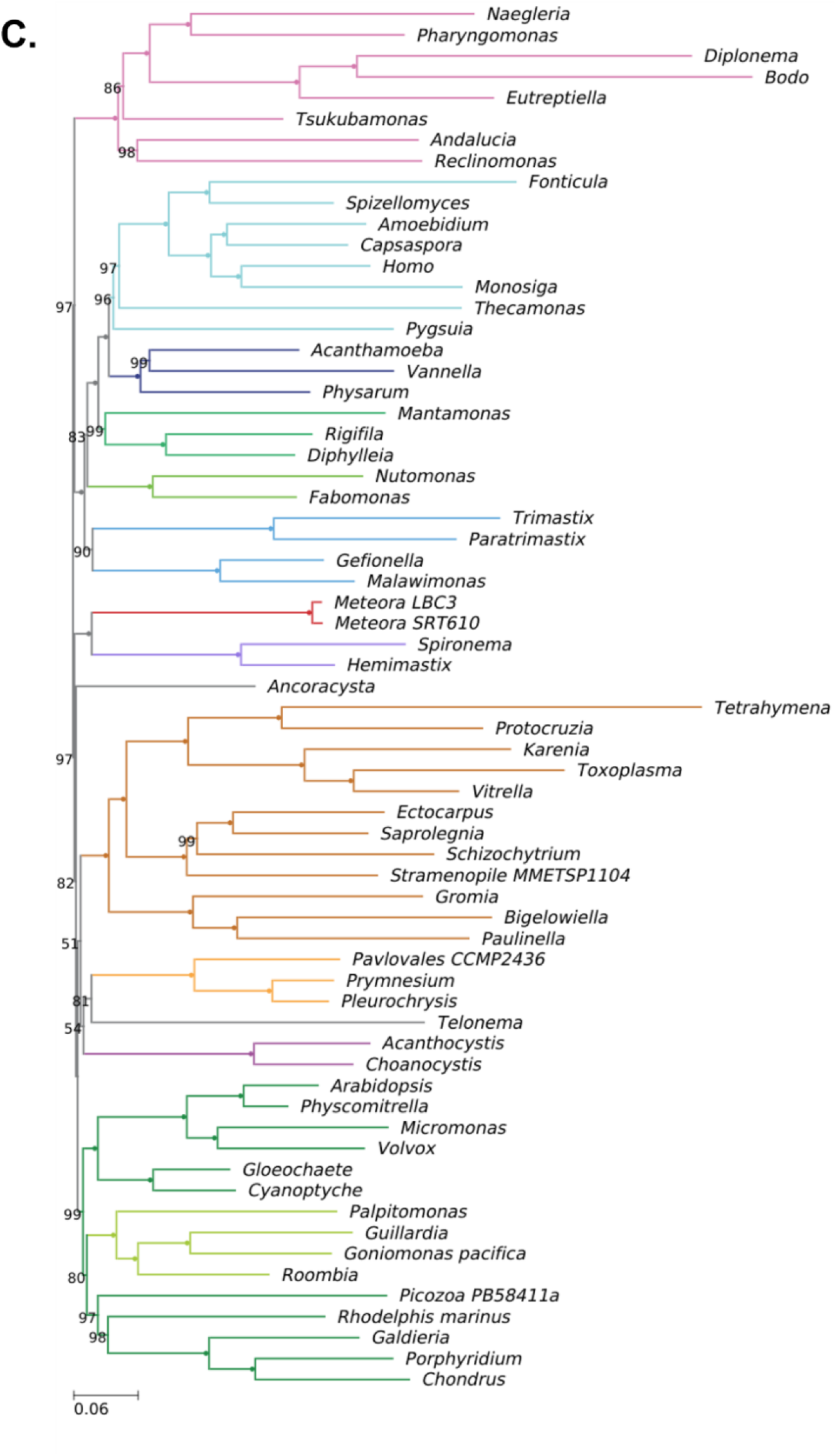
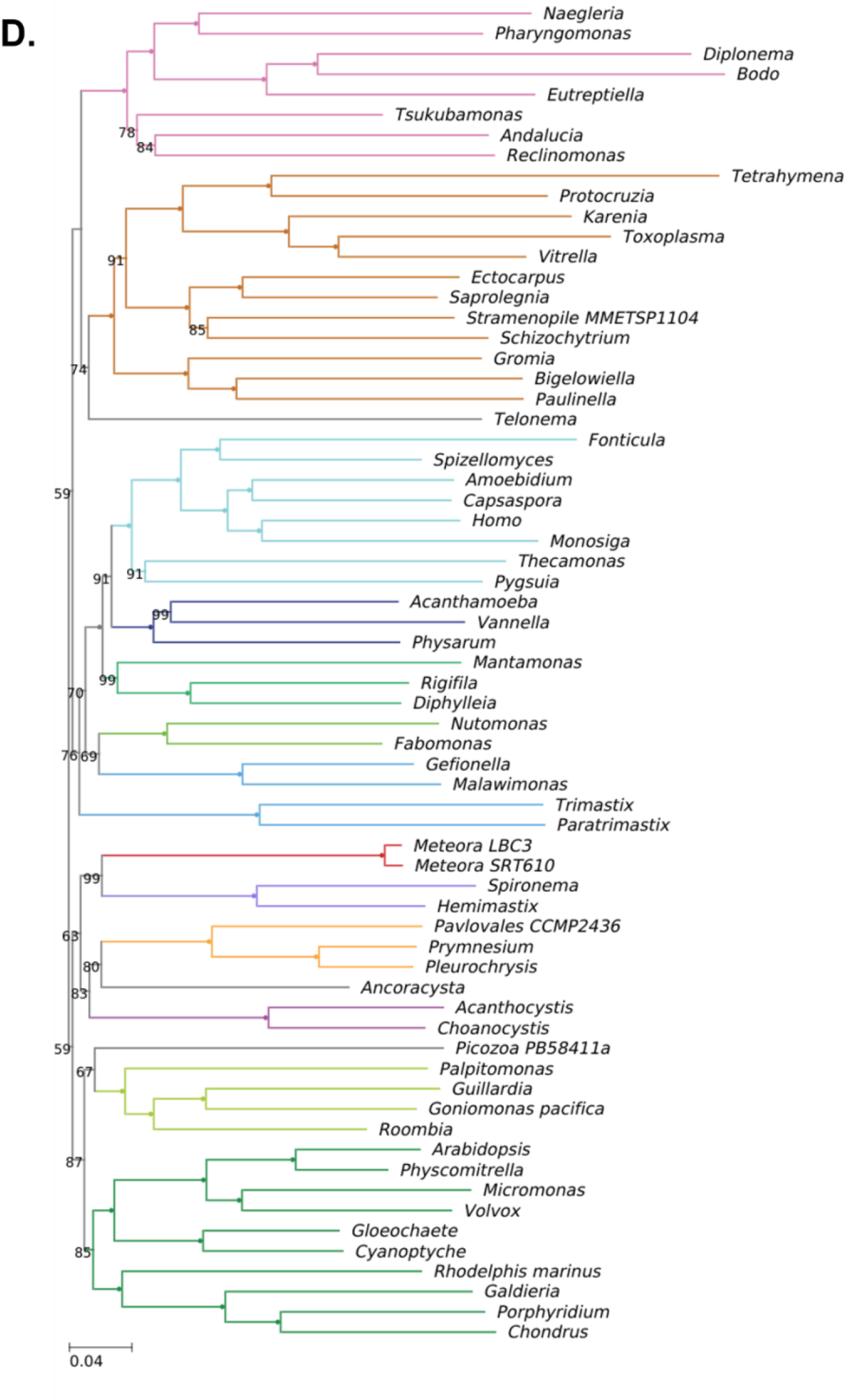
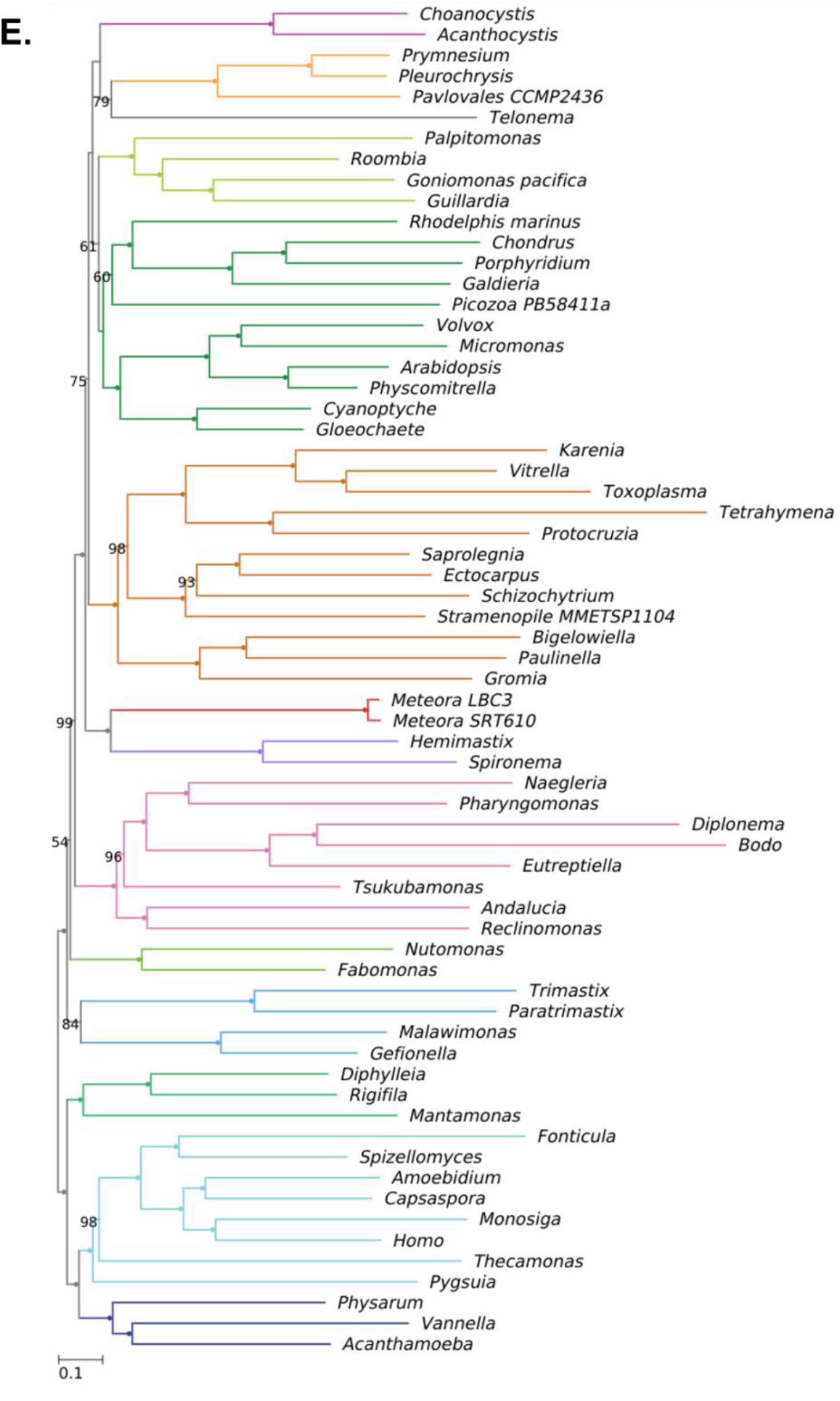
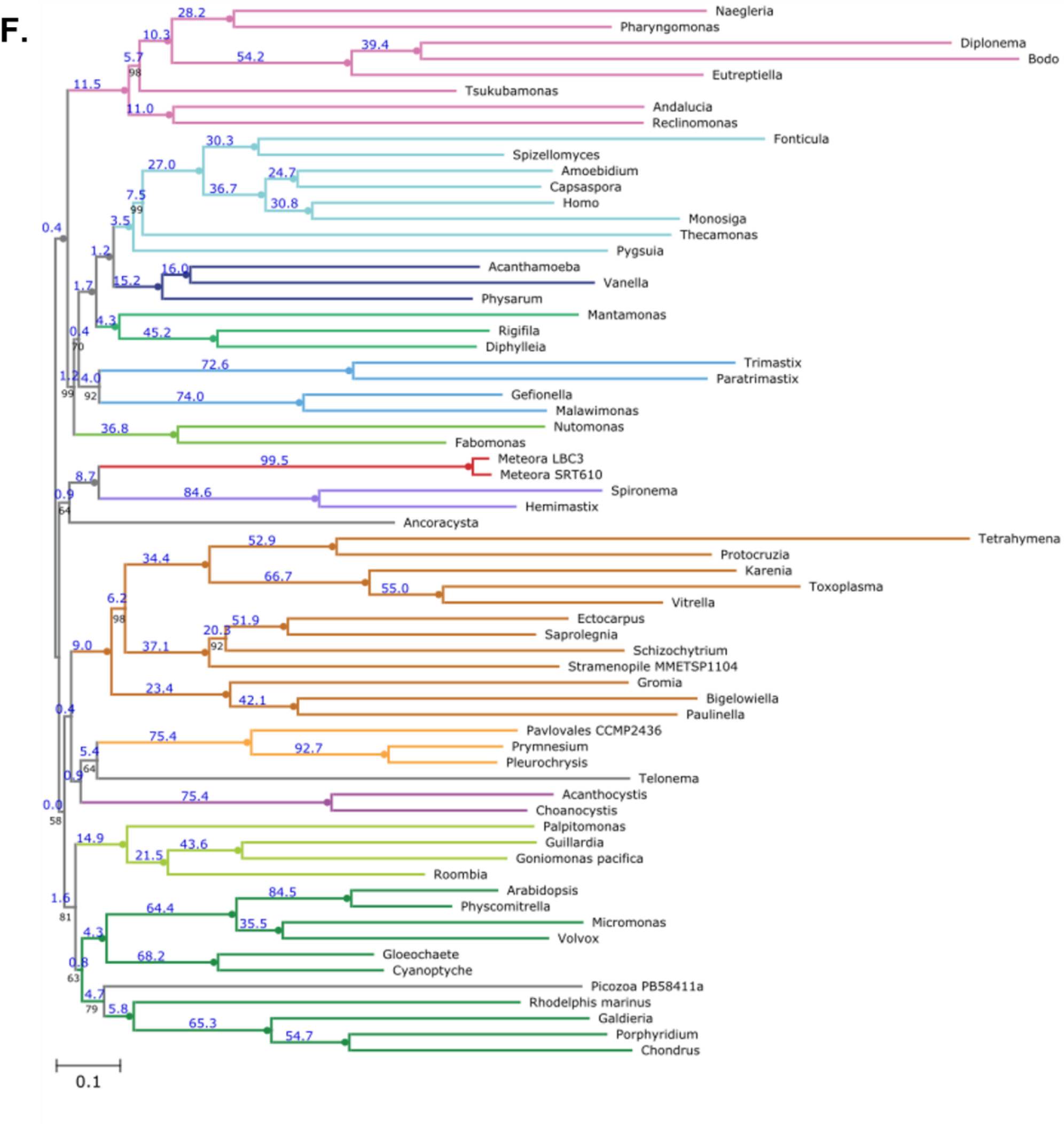
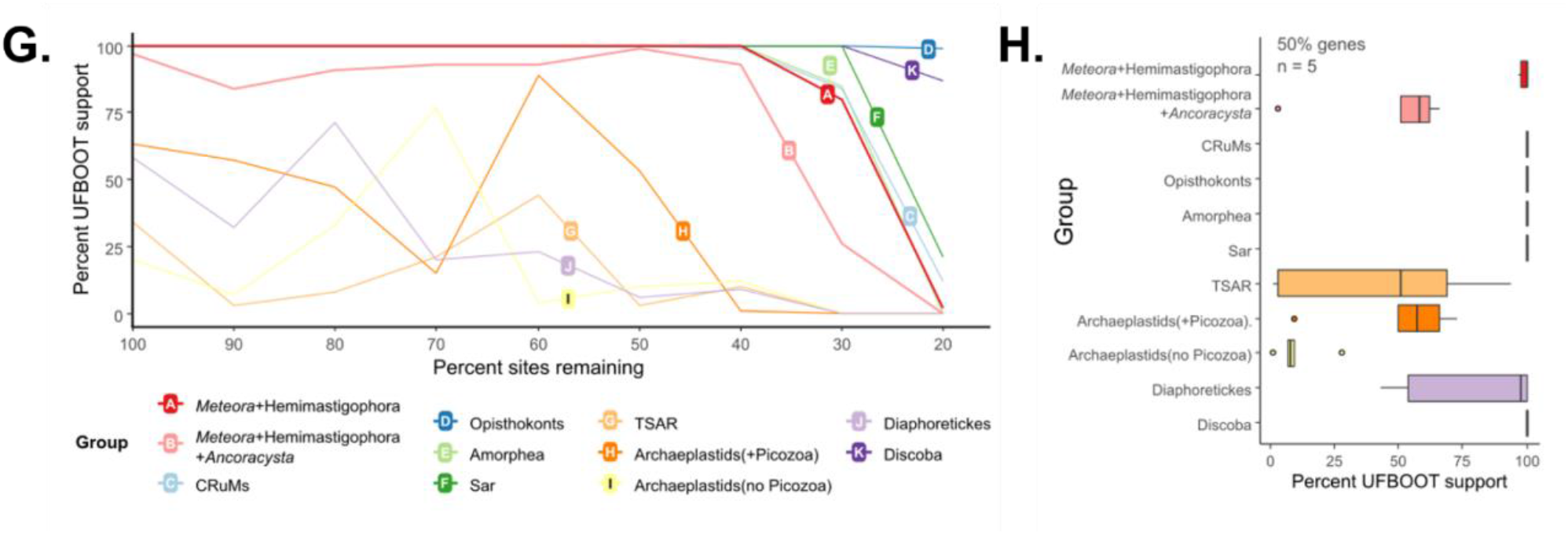
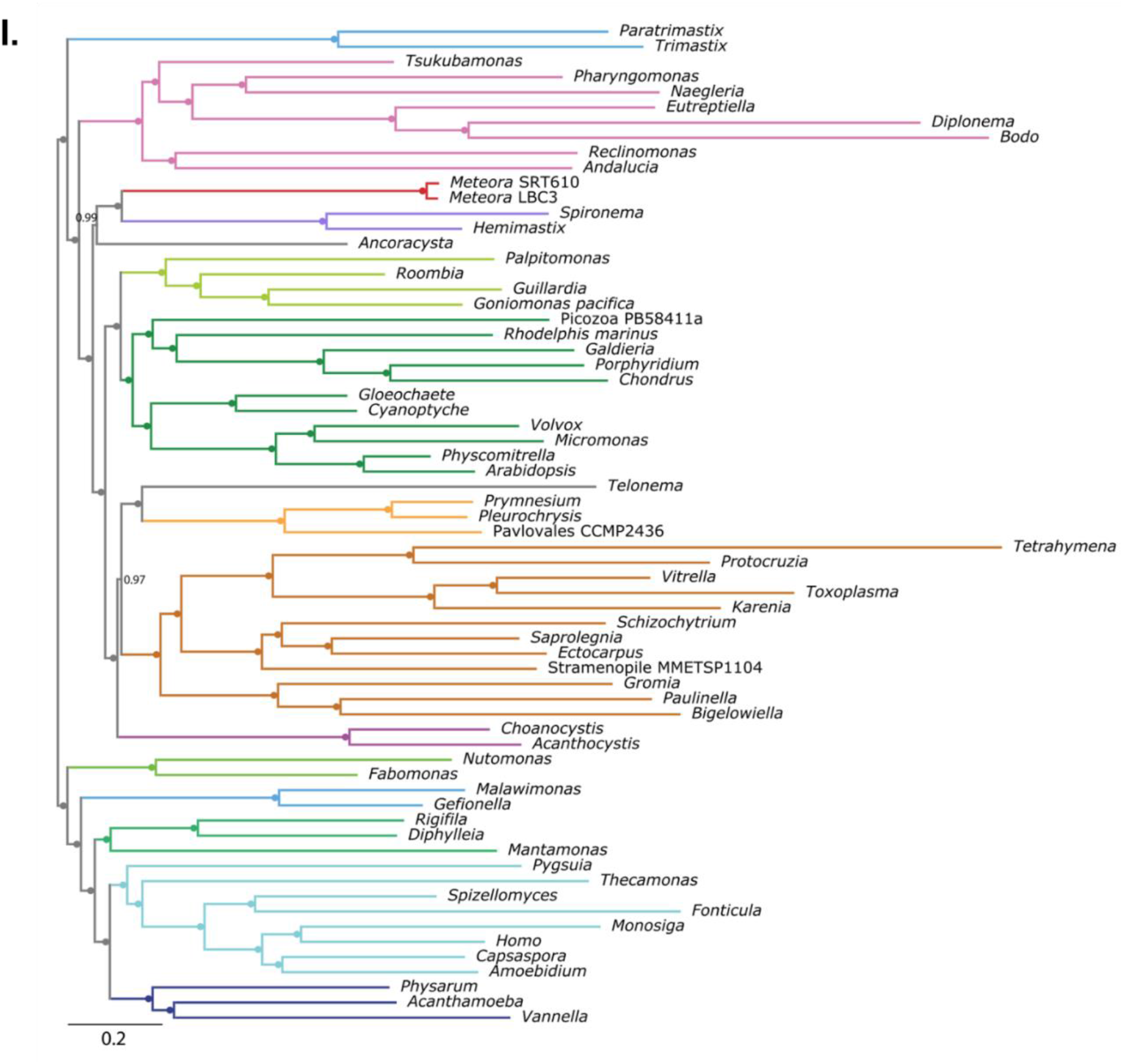
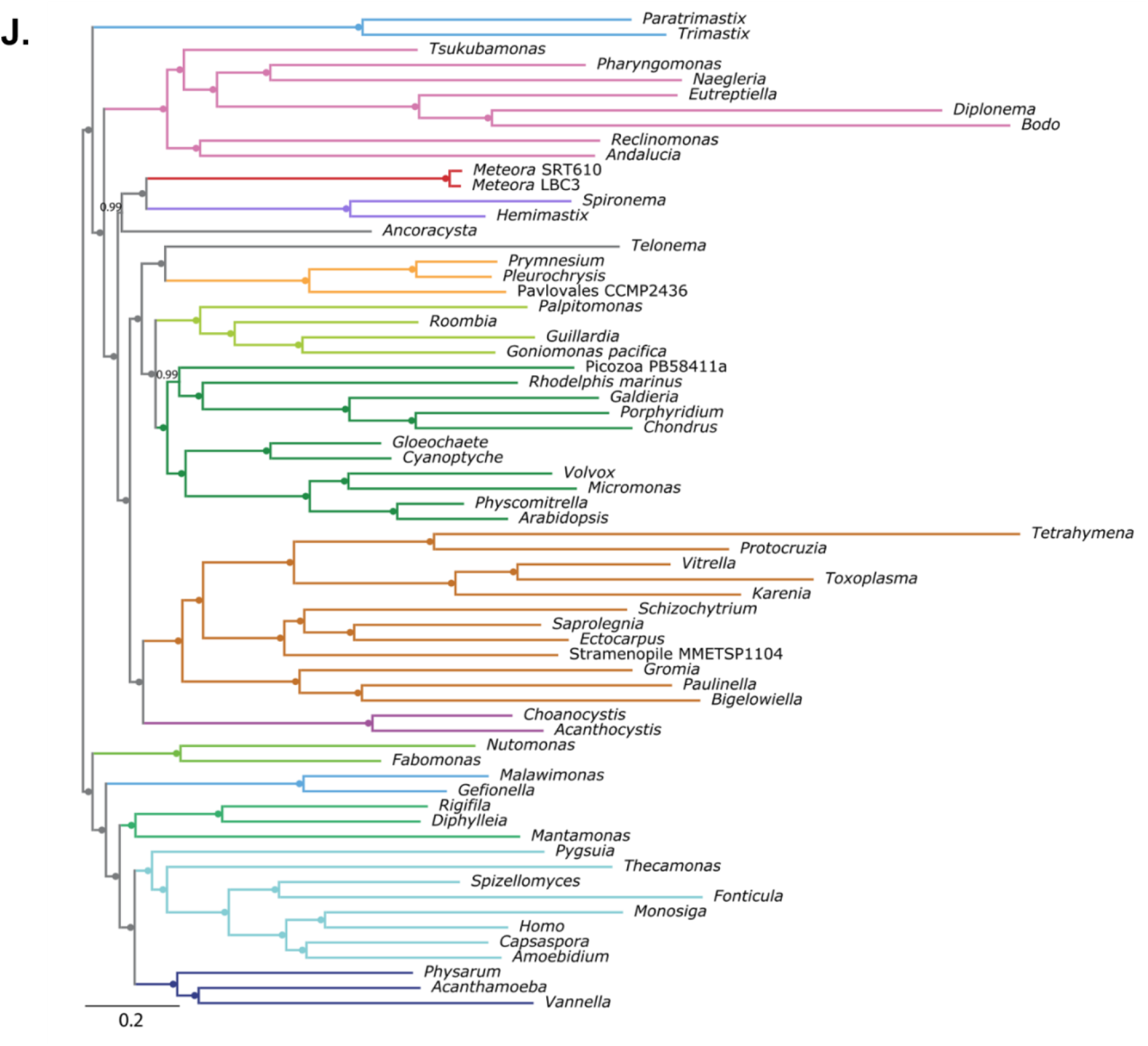
Additional/supplementary phylogenomic analyses. A) 108 taxon phylogeny inferred from 70471 sites across a concatenated 254-gene alignment under the LG+C20+F+Γ model, representing major eukaryotic groups. Node support values represent % UFBOOT support from 1000 replicates. Filled circles indicate full support. Bars on the right represent coverage across the alignment as percent genes (top) and percent sites (bottom). B) No long-branching taxa (nLB) phylogeny, core 66-taxon dataset with three taxa removed, inferred from a concatenated 254-gene alignment under the LG+MAM60+Γ model, support values from 1000 UFBOOT replicates. Filled circles indicate full support. C) Phylogeny inferred from SR4-recoded 254-gene alignment derived from core 66-taxon dataset, under the LG+MAM60+Γ model, support values from 1000 UFBOOT replicates. Filled circles indicate full support. D) Phylogeny inferred from MinMax-Chisq-recoded 254-gene alignment derived from core 66-taxon dataset, under the LG+MAM60+Γ model, support values from 1000 UFBOOT replicates. Filled circles indicate full support. E) No *Ancoracysta* (nAnco) phylogeny; core 66-taxon dataset with *Ancoracysta* removed, inferred from a concatenated 254-gene alignment under the LG+MAM60+Γ model, support values from 1000 UFBOOT replicates. Filled circles indicate full support. F) 66-taxon topology (see Fig 3) with gene concordance factor values indicated on branches as percentages in blue. G) Fast-site removal (FSR) profile of selected groupings with step-wise removal in 10% increments. Plot traces UFBOOT support (1000 replicates) under the LG+MAM60+Γ model. H) Support for selected groupings following 50% gene jackknifing (i.e., 50% of genes randomly removed) across 5 replicates. (trees in datadryad). I) 66-taxon PhyloBayes CAT+GTR consensus phylogeny of chains 2-4, following 1.1 x 10^4^ cycles with a burn-in of 500. Support values show posterior probabilities. Filled circles indicate full support. J) 66-taxon PhyloBayes CAT+GTR phylogeny of chain 1, following 1.1 x 10^4^ cycles with a burn-in of 500. Support values show posterior probabilities. Filled circles indicate full support.

## Bibliography

1. Burki, F., Roger, A.J., Brown, M.W., and Simpson, A.G.B. (2020). The New Tree of Eukaryotes. Trends in Ecology & Evolution 35, 43–55. 10.1016/j.tree.2019.08.008.

2. Lax, G., Eglit, Y., Eme, L., Bertrand, E.M., Roger, A.J., and Simpson, A.G.B. (2018). Hemimastigophora is a novel supra-kingdom-level lineage of eukaryotes. Nature 564, 410–414. 10.1038/s41586-018-0708-8.

3. Tikhonenkov, D.V., Mikhailov, K.V., Hehenberger, E., Karpov, S.A., Prokina, K.I., Esaulov, A.S., Belyakova, O.I., Mazei, Y.A., Mylnikov, A.P., Aleoshin, V.V., et al. (2020). New Lineage of Microbial Predators Adds Complexity to Reconstructing the Evolutionary Origin of Animals. Current Biology. 10.1016/j.cub.2020.08.061.

4. Gawryluk, R.M.R., Tikhonenkov, D.V., Hehenberger, E., Husnik, F., Mylnikov, A.P., and Keeling, P.J. (2019). Non-photosynthetic predators are sister to red algae. Nature 572, 240–243. 10.1038/s41586-019-1398-6.

5. Hausmann, K., Weitere, M., Wolf, M., and Arndt, H. (2002). Meteora sporadica gen. nov. et sp. nov. (Protista incertae sedis) – an extraordinary free-living protist from the Mediterranean deep sea. European Journal of Protistology 38, 171–177. 10.1078/0932-4739-00872.

6. Galindo, L.J., López-García, P., and Moreira, D. (2022). First Molecular Characterization of the Elusive Marine Protist Meteora sporadica. Protist 173, 125896. 10.1016/j.protis.2022.125896.

7. Hausmann, K., Hülsmann, N., and Radek, R. (2003). Protistology. Protistology. Archibald, J.M., Simpson, A.G.B., and Slamovits, C.H. eds. (2017). Handbook of the Protists (Springer International Publishing) 10.1007/978-3-319-28149-0.

8. Mikrjukov, K.A. (1995). Structure, function, and formation of extrusive organellesmicrotoxicysts in the rhizopodPenardia cometa. Protoplasma 188, 186–191. 10.1007/BF01280370.

9. Lax, G., and Simpson, A.G.B. (2020). The Molecular Diversity of Phagotrophic Euglenids Examined Using Single-cell Methods. Protist, 125757. 10.1016/j.protis.2020.125757.

10. Howe, A.T., Bass, D., Vickerman, K., Chao, E.E., and Cavalier-Smith, T. (2009). Phylogeny, taxonomy, and astounding genetic diversity of glissomonadida ord. nov., the dominant gliding zooflagellates in soil (Protozoa: Cercozoa). Protist 160, 159–189. 10.1016/j.protis.2008.11.007.

11. Glücksman, E., Snell, E.A., Berney, C., Chao, E.E., Bass, D., and Cavalier-Smith, T. (2011). The novel marine gliding zooflagellate genus Mantamonas (Mantamonadida ord. n.: Apusozoa). Protist 162, 207–221. 10.1016/j.protis.2010.06.004.

12. Torruella, G., Moreira, D., and López-García, P. (2017). Phylogenetic and ecological diversity of apusomonads, a lineage of deep-branching eukaryotes. Environ Microbiol Rep 9, 113–119. 10.1111/1758-2229.12507.

13. Mahé, F., de Vargas, C., Bass, D., Czech, L., Stamatakis, A., Lara, E., Singer, D., Mayor, J., Bunge, J., Sernaker, S., et al. (2017). Parasites dominate hyperdiverse soil protist communities in Neotropical rainforests. Nat Ecol Evol 1, 1–8. 10.1038/s41559-017-0091.

14. Strassert, J.F.H., Jamy, M., Mylnikov, A.P., Tikhonenkov, D.V., and Burki, F. (2019). New Phylogenomic Analysis of the Enigmatic Phylum Telonemia Further Resolves the Eukaryote Tree of Life. Molecular Biology and Evolution 36, 757–765. 10.1093/molbev/msz012.

15. Tikhonenkov, D.V., Mikhailov, K.V., Gawryluk, R.M.R., Belyaev, A.O., Mathur, V., Karpov, S.A., Zagumyonnyi, D.G., Borodina, A.S., Prokina, K.I., Mylnikov, A.P., et al. (2022). Microbial predators form a new supergroup of eukaryotes. Nature 612, 714–719. 10.1038/s41586-022-05511-5.

16. Susko, E., and Roger, A.J. (2007). On Reduced Amino Acid Alphabets for Phylogenetic Inference. Molecular Biology and Evolution 24, 2139–2150. 10.1093/molbev/msm144.

17. Susko, E. minmax-chisq: Obtaining a Reduced Amino Acid Alphabet with More Homogeneous Composition, Version 1.1.

18. Minh, B.Q., Hahn, M.W., and Lanfear, R. (2020). New Methods to Calculate Concordance Factors for Phylogenomic Datasets. Molecular Biology and Evolution 37, 2727–2733. 10.1093/molbev/msaa106.

19. Janouškovec, J., Tikhonenkov, D.V., Burki, F., Howe, A.T., Rohwer, F.L., Mylnikov, A.P., and Keeling, P.J. (2017). A New Lineage of Eukaryotes Illuminates Early Mitochondrial Genome Reduction. Current Biology 27, 3717–3724.e5. 10.1016/j.cub.2017.10.051.

20. Lang, B.F., Burger, G., O’Kelly, C.J., Cedergren, R., Golding, G.B., Lemieux, C., Sankoff, D., Turmel, M., and Gray, M.W. (1997). An ancestral mitochondrial DNA resembling a eubacterial genome in miniature. Nature 387, 493–497. 10.1038/387493a0.

21. Burger, G., Gray, M.W., Forget, L., and Lang, B.F. (2013). Strikingly Bacteria-Like and Gene-Rich Mitochondrial Genomes throughout Jakobid Protists. Genome Biology and Evolution 5, 418–438. 10.1093/gbe/evt008.

22. Yazaki, E., Yabuki, A., Nishimura, Y., Shiratori, T., Hashimoto, T., and Inagaki, Y. (2022). Microheliella maris possesses the most gene-rich mitochondrial genome in Diaphoretickes. Front. Ecol. Evol. 10, 1030570. 10.3389/fevo.2022.1030570.

23. Kamikawa, R., Shiratori, T., Ishida, K.-I., Miyashita, H., and Roger, A.J. (2016). Group II Intron-Mediated *Trans* -Splicing in the Gene-Rich Mitochondrial Genome of an Enigmatic Eukaryote, *Diphylleia rotans*. Genome Biol Evol 8, 458–466. 10.1093/gbe/evw011.

24. Yazaki, E., Yabuki, A., Imaizumi, A., Kume, K., Hashimoto, T., and Inagaki, Y. (2022). The closest lineage of Archaeplastida is revealed by phylogenomics analyses that include *Microheliella maris*. Open Biol. 12, 210376. 10.1098/rsob.210376.

25. Herman, E.K., Greninger, A.L., Visvesvara, G.S., Marciano-Cabral, F., Dacks, J.B., and Chiu, C.Y. (2013). The Mitochondrial Genome and a 60-kb Nuclear DNA Segment from *Naegleria fowleri*, the Causative Agent of Primary Amoebic Meningoencephalitis. J. Eukaryot. Microbiol. 60, 179–191. 10.1111/jeu.12022.

26. Kamikawa, R., Kolisko, M., Nishimura, Y., Yabuki, A., Brown, M.W., Ishikawa, S.A., Ishida, K., Roger, A.J., Hashimoto, T., and Inagaki, Y. (2014). Gene Content Evolution in Discobid Mitochondria Deduced from the Phylogenetic Position and Complete Mitochondrial Genome of Tsukubamonas globosa. Genome Biology and Evolution 6, 306–315. 10.1093/gbe/evu015.

27. Fu, C.-J., Sheikh, S., Miao, W., Andersson, S.G.E., and Baldauf, S.L. (2014). Missing Genes, Multiple ORFs, and C-to-U Type RNA Editing in Acrasis kona (Heterolobosea, Excavata) Mitochondrial DNA. Genome Biology and Evolution 6, 2240–2257. 10.1093/gbe/evu180.

28. Yang, J., Harding, T., Kamikawa, R., Simpson, A.G.B., and Roger, A.J. (2017). Mitochondrial Genome Evolution and a Novel RNA Editing System in Deep-Branching Heteroloboseids. Genome Biology and Evolution 9, 1161–1174. 10.1093/gbe/evx086.

29. Yabuki, A., Gyaltshen, Y., Heiss, A.A., Fujikura, K., and Kim, E. (2018). Ophirina amphinema n. gen., n. sp., a New Deeply Branching Discobid with Phylogenetic Affinity to Jakobids. Sci Rep 8, 16219. 10.1038/s41598-018-34504-6.

30. Janouškovec, J., Tikhonenkov, D.V., Burki, F., Howe, A.T., Rohwer, F.L., Mylnikov, A.P., and Keeling, P.J. (2017). A New Lineage of Eukaryotes Illuminates Early Mitochondrial Genome Reduction. Current Biology 27, 3717–3724.e5. 10.1016/j.cub.2017.10.051.

31. Nishimura, Y., Shiratori, T., Ishida, K., Hashimoto, T., Ohkuma, M., and Inagaki, Y. (2019). Horizontally-acquired genetic elements in the mitochondrial genome of a centrohelid Marophrys sp. SRT127. Sci Rep 9, 4850. 10.1038/s41598-019-41238-6.

32. Allen, J.W.A., Jackson, A.P., Rigden, D.J., Willis, A.C., Ferguson, S.J., and Ginger, M.L. (2008). Order within a mosaic distribution of mitochondrial c-type cytochrome biogenesis systems?: Evolution of mitochondrial cytochrome c maturation. FEBS Journal 275, 2385– 2402. 10.1111/j.1742-4658.2008.06380.x.

33. Cavalier-Smith, T. (2010). Kingdoms Protozoa and Chromista and the eozoan root of the eukaryotic tree. Biol. Lett. 6, 342–345. 10.1098/rsbl.2009.0948.

34. Foissner, W., Blatterer, H., and Foissner, I. (1988). The hemimastigophora (Hemimastix amphikineta nov. gen., nov. spec.), a new protistan phylum from gondwanian soils. European journal of protistology 23, 361–383. 10.1016/S0932-4739(88)80027-0.

35. Grattepanche, J.-D., Walker, L.M., Ott, B.M., Paim Pinto, D.L., Delwiche, C.F., Lane, C.E., and Katz, L.A. (2018). Microbial Diversity in the Eukaryotic SAR Clade: Illuminating the Darkness Between Morphology and Molecular Data. BioEssays 40, 1700198. 10.1002/bies.201700198.

36. Brown, M.W., Heiss, A.A., Kamikawa, R., Inagaki, Y., Yabuki, A., Tice, A.K., Shiratori, T., Ishida, K.-I., Hashimoto, T., Simpson, A.G.B., et al. (2018). Phylogenomics Places Orphan Protistan Lineages in a Novel Eukaryotic Super-Group. Genome Biology and Evolution 10, 427–433. 10.1093/gbe/evy014.

37. Rasband, W.S. (1997). ImageJ, U. S. National Institutes of Health, Bethesda, Maryland, USA.

38. Schneider, C.A., Rasband, W.S., and Eliceiri, K.W. (2012). NIH Image to ImageJ: 25 years of image analysis. Nat Methods 9, 671–675. 10.1038/nmeth.2089.

39. Gouy, M., Guindon, S., and Gascuel, O. (2010). SeaView version 4: A multiplatform graphical user interface for sequence alignment and phylogenetic tree building. Molecular biology and evolution 27, 221–224. 10.1093/molbev/msp259.

40. Edgar, R.C. (2004). MUSCLE: multiple sequence alignment with high accuracy and high throughput. Nucleic Acids Res 32, 1792–1797. 10.1093/nar/gkh340.

41. Jamy, M., Foster, R., Barbera, P., Czech, L., Kozlov, A., Stamatakis, A., Bending, G., Hilton, S., Bass, D., and Burki, F. (2020). Long-read metabarcoding of the eukaryotic rDNA operon to phylogenetically and taxonomically resolve environmental diversity. Molecular Ecology Resources 20, 429–443. https://doi.org/10.1111/1755-0998.13117.

42. Castresana, J. (2000). Selection of Conserved Blocks from Multiple Alignments for Their Use in Phylogenetic Analysis. Molecular Biology and Evolution 17, 540–552. 10.1093/oxfordjournals.molbev.a026334.

43. Stamatakis, A. (2014). RAxML version 8: a tool for phylogenetic analysis and post-analysis of large phylogenies. Bioinformatics (Oxford, England) 30, 1312–1313. 10.1093/bioinformatics/btu033.

44. Seeman, T. (2018). Bacterial ribosomal RNA predictor.

45. de Vargas, C., Audic, S., Henry, N., Decelle, J., Mahé, F., Logares, R., Lara, E., Berney, C., Le Bescot, N., Probert, I., et al. (2015). Ocean plankton. Eukaryotic plankton diversity in the sunlit ocean. Science (New York, N.Y.) 348, 1261605–1261605. 10.1126/science.1261605.

46. Huse, S.M., Mark Welch, D.B., Voorhis, A., Shipunova, A., Morrison, H.G., Eren, A.M., and Sogin, M.L. (2014). VAMPS: a website for visualization and analysis of microbial population structures. BMC Bioinformatics 15, 41. 10.1186/1471-2105-15-41.

47. Pawlowski, J., Audic, S., Adl, S., Bass, D., Belbahri, L., Berney, C., Bowser, S.S., Cepicka, I., Decelle, J., Dunthorn, M., et al. (2012). CBOL Protist Working Group: Barcoding Eukaryotic Richness beyond the Animal, Plant, and Fungal Kingdoms. PLOS Biology 10, e1001419. 10.1371/journal.pbio.1001419.

48. Vaulot, D., Hung Sim, C.W., Ong, D., Teo, B., Biwer, C., Jamy, M., and dos Santos, A.L. (2022). metaPR2: a database of eukaryotic 18S rRNA metabarcodes with an emphasis on protists. bioRxiv, 2022.02.04.479133. 10.1101/2022.02.04.479133.

49. Schoenle, A., Hohlfeld, M., Hermanns, K., Mahé, F., de Vargas, C., Nitsche, F., and Arndt, H. (2021). High and specific diversity of protists in the deep-sea basins dominated by diplonemids, kinetoplastids, ciliates and foraminiferans. Commun Biol 4, 1–10. 10.1038/s42003-021-02012-5.

50. Obiol, A., Giner, C.R., Sánchez, P., Duarte, C.M., Acinas, S.G., and Massana, R. (2020). A metagenomic assessment of microbial eukaryotic diversity in the global ocean. Molecular Ecology Resources 20, 718–731. 10.1111/1755-0998.13147.

51. Geisen, S., Tveit, A.T., Clark, I.M., Richter, A., Svenning, M.M., Bonkowski, M., and Urich, T. (2015). Metatranscriptomic census of active protists in soils. ISME J 9, 2178–2190. 10.1038/ismej.2015.30.

52. Suter, E.A., Pachiadaki, M., Taylor, G.T., and Edgcomb, V.P. (2021). Eukaryotic Parasites Are Integral to a Productive Microbial Food Web in Oxygen-Depleted Waters. Front Microbiol 12, 764605. 10.3389/fmicb.2021.764605.

53. Berger, S., and Stamatakis, A. (2012). PaPaRa 2. 0 : A Vectorized Algorithm for Probabilistic Phylogeny-Aware Alignment Extension. https://cme.h-its.org/exelixis/pubs/Exelixis-RRDR-2012-5.pdf.

54. Berger, S.A., Krompass, D., and Stamatakis, A. (2011). Performance, Accuracy, and Web Server for Evolutionary Placement of Short Sequence Reads under Maximum Likelihood. Systematic Biology 60, 291–302. 10.1093/sysbio/syr010.

55. Yu, G., Smith, D.K., Zhu, H., Guan, Y., and Lam, T.T.-Y. (2017). ggtree: an r package for visualization and annotation of phylogenetic trees with their covariates and other associated data. Methods in Ecology and Evolution 8, 28–36. 10.1111/2041-210X.12628.

56. Andrews, S. (2010). FastQC: A Quality Control Tool for High Throughput Sequence Data [Online].

57. Bolger, A.M., Lohse, M., and Usadel, B. (2014). Trimmomatic: a flexible trimmer for Illumina sequence data. Bioinformatics (Oxford, England) 30, 2114–2120. 10.1093/bioinformatics/btu170.

58. Haas, B.J., Papanicolaou, A., Yassour, M., Grabherr, M., Blood, P.D., Bowden, J., Couger, M.B., Eccles, D., Li, B., Lieber, M., et al. (2013). De novo transcript sequence reconstruction from RNA-seq using the Trinity platform for reference generation and analysis. Nature protocols 8, 1494–1512. 10.1038/nprot.2013.084.

59. Simão, F.A., Waterhouse, R.M., Ioannidis, P., Kriventseva, E.V., and Zdobnov, E.M. (2015). BUSCO: assessing genome assembly and annotation completeness with single-copy orthologs. Bioinformatics 31, 3210–3212. 10.1093/bioinformatics/btv351.

60. Katoh, K., and Standley, D.M. (2013). MAFFT multiple sequence alignment software version 7: improvements in performance and usability. Molecular biology and evolution 30, 772–780. 10.1093/molbev/mst010.

61. Criscuolo, A., and Gribaldo, S. (2010). BMGE (Block Mapping and Gathering with Entropy): a new software for selection of phylogenetic informative regions from multiple sequence alignments. BMC evolutionary biology 10, 210–210. 10.1186/1471-2148-10-210.

62. Le, S.Q., Dang, C.C., and Gascuel, O. (2012). Modeling Protein Evolution with Several Amino Acid Replacement Matrices Depending on Site Rates. Molecular Biology and Evolution 29, 2921–2936. 10.1093/molbev/mss112.

63. Nguyen, L.-T., Schmidt, H.A., von Haeseler, A., and Minh, B.Q. (2015). IQ-TREE: a fast and effective stochastic algorithm for estimating maximum-likelihood phylogenies. Molecular biology and evolution 32, 268–274. 10.1093/molbev/msu300.

64. Whelan, S., Irisarri, I., and Burki, F. (2018). PREQUAL: detecting non-homologous characters in sets of unaligned homologous sequences. Bioinformatics 34, 3929–3930. 10.1093/bioinformatics/bty448.

65. Minh, B.Q., Nguyen, M.A.T., and von Haeseler, A. (2013). Ultrafast approximation for phylogenetic bootstrap. Molecular biology and evolution 30, 1188–1195. 10.1093/molbev/mst024.

66. Susko, E., Lincker, L., and Roger, A.J. (2018). Accelerated Estimation of Frequency Classes in Site-Heterogeneous Profile Mixture Models. Molecular Biology and Evolution 35, 1266–1283. 10.1093/molbev/msy026.

67. Susko, E. (2022). MAMMaL: (M)ultinomial (A)pproximate (M)ixture (Ma)ximum (L)ikelihood Accelerated Estimation of Frequency Classes in Site-heterogeneous Profile Mixture Models Version 1.1.3.

68. Wang, H.-C., Minh, B.Q., Susko, E., and Roger, A.J. (2018). Modeling Site Heterogeneity with Posterior Mean Site Frequency Profiles Accelerates Accurate Phylogenomic Estimation. Syst Biol 67, 216–235. 10.1093/sysbio/syx068.

69. Lartillot, N., and Philippe, H. (2004). A Bayesian Mixture Model for Across-Site Heterogeneities in the Amino-Acid Replacement Process. Molecular Biology and Evolution 21, 1095–1109. 10.1093/molbev/msh112.

70. Tice, A.K., Žihala, D., Pánek, T., Jones, R.E., Salomaki, E.D., Nenarokov, S., Burki, F., Eliáš, M., Eme, L., Roger, A.J., et al. (2021). PhyloFisher: A phylogenomic package for resolving eukaryotic relationships. PLOS Biology 19, e3001365. 10.1371/journal.pbio.3001365.

71. Minh, B.Q., Schmidt, H.A., Chernomor, O., Schrempf, D., Woodhams, M.D., von Haeseler, A., and Lanfear, R. (2020). IQ-TREE 2: New Models and Efficient Methods for Phylogenetic Inference in the Genomic Era. Molecular Biology and Evolution 37, 1530–1534. 10.1093/molbev/msaa015.

72. Huerta-Cepas, J., Serra, F., and Bork, P. (2016). ETE 3: Reconstruction, Analysis, and Visualization of Phylogenomic Data. Molecular Biology and Evolution 33, 1635–1638. 10.1093/molbev/msw046.

73. Keller, M.D., Selvin, R.C., Claus, W., and Guillard, R.R.L. (1987). MEDIA FOR THE CULTURE OF OCEANIC ULTRAPHYTOPLANKTON1,2. Journal of Phycology 23, 633–638. 10.1111/j.1529-8817.1987.tb04217.x.

74. Bolger, A.M., Lohse, M., and Usadel, B. (2014). Trimmomatic: a flexible trimmer for Illumina sequence data. Bioinformatics 30, 2114–2120. 10.1093/bioinformatics/btu170.

75. Langmead, B., and Salzberg, S.L. (2012). Fast gapped-read alignment with Bowtie 2. Nat Methods 9, 357–359. 10.1038/nmeth.1923.

76. Lin, Y., Yuan, J., Kolmogorov, M., Shen, M.W., Chaisson, M., and Pevzner, P.A. (2016). Assembly of long error-prone reads using de Bruijn graphs. Proc Natl Acad Sci USA 113, E8396–E8405. 10.1073/pnas.1604560113.

77. Kolmogorov, M., Yuan, J., Lin, Y., and Pevzner, P.A. (2019). Assembly of long, error-prone reads using repeat graphs. Nat Biotechnol 37, 540–546. 10.1038/s41587-019-0072-8.

78. Kolmogorov, M., Bickhart, D.M., Behsaz, B., Gurevich, A., Rayko, M., Shin, S.B., Kuhn, K., Yuan, J., Polevikov, E., Smith, T.P.L., et al. (2020). metaFlye: scalable long-read metagenome assembly using repeat graphs. Nat Methods 17, 1103–1110. 10.1038/s41592-020-00971-x.

79. Kim, D., Paggi, J.M., Park, C., Bennett, C., and Salzberg, S.L. (2019). Graph-based genome alignment and genotyping with HISAT2 and HISAT-genotype. Nat Biotechnol 37, 907–915. 10.1038/s41587-019-0201-4.

80. Sedlazeck, F.J., Rescheneder, P., Smolka, M., Fang, H., Nattestad, M., von Haeseler, A., and Schatz, M.C. (2018). Accurate detection of complex structural variations using single-molecule sequencing. Nat Methods 15, 461–468. 10.1038/s41592-018-0001-7.

81. Walker, B.J., Abeel, T., Shea, T., Priest, M., Abouelliel, A., Sakthikumar, S., Cuomo, C.A., Zeng, Q., Wortman, J., Young, S.K., et al. (2014). Pilon: An Integrated Tool for Comprehensive Microbial Variant Detection and Genome Assembly Improvement. PLoS ONE 9, e112963. 10.1371/journal.pone.0112963.

82. Milne, I., Stephen, G., Bayer, M., Cock, P.J.A., Pritchard, L., Cardle, L., Shaw, P.D., and Marshall, D. (2013). Using Tablet for visual exploration of second-generation sequencing data. Briefings in Bioinformatics 14, 193–202. 10.1093/bib/bbs012.

83. Derelle, R., Torruella, G., Klimeš, V., Brinkmann, H., Kim, E., Vlček, Č., Lang, B.F., and Eliáš, M. (2015). Bacterial proteins pinpoint a single eukaryotic root. Proc. Natl. Acad. Sci. U.S.A. 112. 10.1073/pnas.1420657112.

